# DLGAP1-AS2-Mediated Phosphatidic Acid Synthesis Confers Chemoresistance via Activation of YAP Signaling

**DOI:** 10.1101/2022.02.24.481869

**Authors:** Yabing Nan, Qingyu Luo, Xiaowei Wu, Shi Liu, Pengfei Zhao, Wan Chang, Aiping Zhou, Zhihua Liu

## Abstract

Squamous cell carcinomas (SCCs) constitute a group of human malignancies that originate from the squamous epithelium. Most SCC patients experience treatment failure and relapse and have a poor prognosis due to de novo and acquired resistance to first-line chemotherapeutic agents. To identify chemoresistance mechanisms and explore novel chemosensitizer targets, we performed whole-transcriptome sequencing of paired resistant/parental SCC cells. We identified DLGAP1 antisense RNA 2 (D-AS2) as a crucial noncoding RNA that contributes to chemoresistance in SCC. Mechanistically, D-AS2 associates with histones to regulate the distal elements of FAM3 metabolism regulating signaling molecule D (FAM3D) and reduces extracellular FAM3D protein secretion. FAM3D interacts with Gα_i_-coupled G protein-coupled receptor (GPCR) formyl peptide receptor (FPR) 1 and FPR2 to suppress phospholipase D (PLD) activity; thus, reduced FAM3D activates PLD signaling. Moreover, activated PLD promotes phosphatidic acid (PA) production and subsequent yes-associated protein (YAP) nuclear translocation. Accordingly, in vivo administration of a D-AS2-targeting antisense oligonucleotide sensitizes SCC to cisplatin treatment. In summary, our study reveals that D-AS2/FAM3D-mediated PLD/PA lipid signaling is essential in SCC chemoresistance and that D-AS2 can be targeted to sensitize SCC to cytotoxic chemotherapeutic agents.

**Significance:** This study identifies D-AS2 as a targetable lipid-related lncRNA that activates YAP signaling via PLD/PA axis to trigger chemoresistance in SCC.

## Introduction

In recent decades, cytotoxic chemotherapeutic regimens including platinum drugs have been widely used for the treatment of human malignancies, with particular benefit in terminal patients who are ineligible for surgery. Unfortunately, chemoresistance can develop via intrinsic or adaptive mechanisms driven by genomic or nongenomic causes. Increasing the sensitivity of cancer cells to chemotherapeutic agents or using chemotherapeutic agents with different targets are the two strategies commonly used to reduce the adverse effects of cytotoxic drugs (1). Encouraging novel targeted therapies have recently emerged in recent years, for example, those targeting CDK4/6, PDL1, and PARP (2–6). However, these targeted therapies exhibit promising effects only in a limited number of subtypes, exhibiting no efficacy in other subtypes. Even for sensitive subtypes, accumulating evidence shows that the combination of targeted therapies and traditional cytotoxic chemotherapeutic agents achieves a better outcome (7, 8). For other subtypes that are not suitable for novel targeted therapies, traditional cytotoxic chemotherapeutic agents remain the first-line treatment. Many attempts have been made to sensitize cancer cells to chemotherapeutic agents or to restore chemoresistance (9–12). However, the mechanisms underlying chemoresistance remain largely undefined. Defining previously unknown resistance mechanisms will identify novel targets to overcome chemoresistance, especially in cancer types whose treatment relies mainly on traditional chemotherapeutic agents.

Metabolic reprogramming in cancer cells has been recognized as an essential process during cancer initiation and progression. However, an increasing number of studies have indicated that cancer metabolic reprogramming is a context-dependent process instead of a unified phenomenon. For example, the “Warburg effect” identified nearly 100 years ago has been challenged by several recent studies that proposed a critical role of oxidative phosphorylation (OXPHOS) in cancer cell survival (13–16). In addition, although increased lipid uptake, lipid storage and lipogenesis usually occur in a variety of cancers and contribute to rapid tumor growth (17–19), some studies have revealed that several lipid and lipid categories, such as palmitic acid and alkyl phospholipids, show anticancer effects (20, 21). As members of a ubiquitous class of lipid metabolism-associated enzymes, phospholipase D (PLD) 1 and PLD2 catalyze the hydrolysis of phosphatidylcholine (PC) into phosphatidic acid (PA) and choline. In addition to PLD-mediated PC hydrolysis, PA can be generated by two other metabolic pathways: lysophosphatidic acid acyltransferase (LPAAT) catalyzes the conversion of lysophosphatidic acid (LPA) to PA, and diacylglycerol kinase (DGK) phosphorylates DAG to produce PA (22). Interestingly, only PA produced via PLD can promote oncogenic Yes-associated protein (YAP) signaling by regulating large tumor suppressor kinase 1 (LATS1) (22, 23). Although activation of YAP signaling is involved in chemoresistance in multiple cancers via TEAD-mediated transcriptional regulation (24–26), the specific mechanism by which YAP mediates chemoresistance in SCC remains unclear. Moreover, how YAP signaling is regulated during the occurrence of chemoresistance requires further exploration.

Squamous cell carcinoma (SCC) originates from the squamous epithelium of various organs (27). The most common locations of SCC include the skin, head and neck, esophagus, lung, and cervix (28). Although SCCs originate from different organs, they share common squamous differentiation markers and genetic mutations (28). The representative traditional cytotoxic drug platinum is still used as the first-line chemotherapy for several SCCs. However, SCC patients usually have a poor prognosis owing to therapeutic resistance and relapse. Thus, SCC is a typical research model for defining chemoresistance mechanisms, and the identified novel therapeutic targets will in turn benefit SCC patients. Herein, we performed whole-transcriptome sequencing of two pairs of chemoresistant and parental SCC cell lines. We identified DLGAP1 antisense RNA 2 (D-AS2) as a critical lipid-related lncRNA that induces chemoresistance. Mechanistically, the direct interaction between D-AS2 and histones blocks the association between H3K27ac and the FAM3 metabolism regulating signaling molecule D (FAM3D) enhancer region and then inhibits the transcription of FAM3D. Low expression of FAM3D activates PLD through the Gα_i_ protein-coupled receptors FPR1 and FPR2, increasing PLD-catalyzed PA production. Furthermore, PA contributes to YAP nuclear translocation and activates nuclear YAP activity.

## Materials and Methods

### Cell Culture

Human esophageal SCC cell lines (29) were kindly provided by Dr Yutaka Shimada (Kyoto University, Japan). The 293T cell line was purchased from the American Type Culture Collection (ATCC; VA, USA). The epidermoid carcinoma cell line A431, tongue SCC cell line CAL-27, and lung SCC cell lines NCI-H1703 and NCI-H596 were purchased from the National Infrastructure of Cell Line Resource (Beijing, China). The tongue SCC cell line UM1 was purchased from Beijing Xigong Biological Technology (Beijing, China). CAL-27 and A431 cells were cultured in DMEM supplemented with 10% fetal bovine serum (FBS). UM1 cells were cultured in DMEM:F12 supplemented with 10% FBS. Other SCC cell lines were cultured in RPMI-1640 medium supplemented with 10% FBS. All cells were maintained in a humidified cell incubator with 5% CO_2_ at 37 °C. All cell lines were routinely verified using short tandem repeat DNA fingerprinting and were tested for *Mycoplasma* contamination using a MycoBlue Mycoplasma Detector (D101; Vazyme Biotech, Nanjing, China) before use in any experiments.

### Patient Samples and In Situ Hybridization (ISH)

Paraffin-embedded esophageal (HEsoS180Su09) and lung (HLugS180Su02) SCC tissue arrays were purchased from Shanghai Outdo Biotechnology (Shanghai, China). Written informed consent was obtained from all patients. ISH assays were performed with an Enhanced Sensitive ISH Detection Kit I (POD) (MK1030; Boster, Wuhan, China) according to the manufacturer’s instructions. In brief, SCC tissue blocks were prehybridized at 40 °C for 3 h and then hybridized at 60 °C overnight with a digoxin-labeled D-AS2 probe. Diaminobenzidine was used as a chromogen, and sections were counterstained with hematoxylin. All slides were scanned using an Aperio scanning system (Aperio, San Diego, CA, USA). H-scores were calculated according to the percentage of positive cells and the staining intensity as follows: H-score = Σpi × i, where pi represents the percentage of positive cells (0–100%) and i represents the staining intensity (0, negative; 1, weak; 2, medium; and 3, strong). ISH staining was scored by three independent observers. The sequences of the probes used for ISH are shown in Supplementary Table S1.

### Xenograft Experiments

All animal protocols were approved by the Animal Care and Use Committee of the Chinese Academy of Medical Sciences Cancer Hospital. For subcutaneous xenografting, 1–5 × 10^6^ SCC cells were injected subcutaneously into 6-week-old male BALB/c nude mice (HFK Bioscience, Beijing, China). NOD-Prkdc^scid^-Il2rg^em1IDMO^ (NPI) mice (IDMO, Beijing, China) were used as hosts for patient-derived xenograft (PDX) model establishment. The experimental details were described previously (30). DDP (5 mg/kg) was administered every 3 days beginning 1–2 weeks after cell implantation. The in vivo-optimized antisense oligonucleotides (ASOs) (RiboBio, Guangzhou, China) were administered by intratumoral injection (5 nmol per injection every 3 days, dissolved in 50 μL of sterile PBS). The tumor volume was calculated as 0.52 × length × width^2^.

### Antibodies and Reagents

The following antibodies were used for western blotting: anti-H1.3 (ab24174, Abcam, MA, USA, 1:1000), anti-H1.5 (ab18208, Abcam, 1:1000), anti-mH2A1 (ab37264, Abcam, 1:1000), anti-H2B (ab1790, Abcam, 1:1000), anti-H3 (ab1791, Abcam, 1:1000), anti-H4 (ab10158, Abcam, 1:1000), anti-YAP (14074, Cell Signaling Technology, MA, USA, 1:1000), anti-p-YAP (S127) (4911, Cell Signaling Technology, 1:1000), anti-LATS1 (3477, Cell Signaling Technology, 1:1000), anti-p-LATS1 (S909) (9157, Cell Signaling Technology, 1:1000), anti-H3K4me1 (5326, Cell Signaling Technology, 1:1000), anti-H3K27me3 (9733, Cell Signaling Technology, 1:1000), anti-H3K9ac (9649, Cell Signaling Technology, 1:1000), anti-H3K27ac (8173, Cell Signaling Technology, 1:1000), anti-H4K16ac (13534, Cell Signaling Technology, 1:1000), and anti-β-actin (A5316, Sigma-Aldrich, MO, USA, 1:4000). The following antibodies were used for RNA-binding protein immunoprecipitation (RIP) assays: anti-H1.3 (ab24174, Abcam, 5 µg/reaction), anti-H1.5 (ab18208, Abcam, 5 μg/reaction), anti-mH2A1 (ab37264, Abcam, 5 μg/reaction), anti-H2B (ab1790, Abcam, 5 μg/reaction), anti-H3 (ab1791, Abcam, 5 μg/reaction), anti-H4 (ab10158, Abcam, 5 μg/reaction), and normal rabbit IgG (PP64B, Millipore, MA, USA; 5 μg/reaction). The following antibodies were used for cleavage under targets & release using nuclease (CUT&RUN) assays: anti-H3K27ac (8173, Cell Signaling Technology, 1:100) and anti-H3K4me1 (5326, Cell Signaling Technology, 1:100). FPR Agonist 43 (HY-19574), HCH6-1 (HY-101283), PBP10 TFA (HY-P1116A), cyclosporine H (HY-P1122), WRW4 (HY-P1119), and R59-022 (HY-107613) were obtained from MedChemExpress (MCE; NJ, USA). CI 976 (C3743) was purchased from Sigma-Aldrich. FIPI (939055-18-2) and CAY10594 (1130067-34-3) were purchased from Topscience (Topscience; Shanghai, China).

### Whole-Transcriptome Sequencing

Samples for each group were collected from two independent cell cultures, and whole-transcriptome sequencing was performed by RiboBio. RNA integrity was evaluated using an Agilent 2200 TapeStation system (Agilent Technologies, CA, USA), and rRNA was removed using an Epicenter Ribo-Zero rRNA Removal Kit (Illumina, CA, USA). After fragmentation into pieces of approximately 200 bp, the purified RNAs were subjected to first-strand and second-strand cDNA synthesis, followed by adaptor ligation and enrichment with a low number of cycles, according to the instructions of an NEBNext Ultra RNA Library Prep Kit for Illumina (New England Biolabs, MA, USA). The purified library products were evaluated using the Agilent 2200 TapeStation system and Qubit 2.0 software (Thermo Scientific, MA, USA) and were then subjected to paired-end sequencing on the Illumina HiSeq 3000 platform (Illumina). Clean reads were obtained after the removal of reads containing adapter or poly-N sequences and low-quality reads. HISAT2 was used to align the clean reads to the reference genome. Differential expression was assessed using DESeq with the read count as input. The Benjamini–Hochberg multiple testing procedure was used, and a change threshold of >2 and an adjusted *P* value of <0.05 were used to identify differentially expressed genes. All differentially expressed genes were used for Kyoto Encyclopedia of Genes and Genomes (KEGG) enrichment analysis. The whole-transcriptome sequencing data can be accessed in the Gene Expression Omnibus database under accession number GSE169337.

### RNA Sequencing (RNA-seq)

The RNA-seq experiment and data analysis were conducted by Shanghai Applied Protein Technology (Shanghai, China). Six replicates per group were prepared from independent cell cultures. The paired-end libraries were prepared using a NEBNext Ultra RNA Library Prep Kit for Illumina (New England Biolabs) following the manufacturer’s instructions, and sequencing was performed using a NovaSeq 6000 instrument (Illumina). A fold change of >2 and *P* value of <0.05 were set as the criteria for identifying significantly differentially expressed genes. The RNA-seq data can be accessed in the Gene Expression Omnibus database under accession number GSE169633.

### Assay for Transposase-Accessible Chromatin with Sequencing (ATAC-Seq)

The ATAC-Seq experiment and data analysis were conducted by Seqhealth Technology (Wuhan, China). In brief, 10 000–50 000 cells were treated with cell lysis buffer, and nuclei were collected by centrifugation for 5 min at 500 × *g*. Tagmentation was performed, and a high-throughput DNA sequencing library was prepared using a TruePrep DNA Library Prep Kit V2 for Illumina (TD501; Vazyme Biotech). The library products were then enriched, quantified, and sequenced on a NovaSeq 6000 sequencer (Illumina) by using the PE150 strategy. Raw data were filtered using Trimmomatic (version 0.36) and deduplicated with FastUniq (version 1.1). Clean reads were mapped to the reference genome by using Bowtie2 software (version 2.2.6) with default parameters. RSeQC software (version 2.6) was used for read distribution analysis, and the insert length was calculated using Collect Insert Size Metrics tools in Picard software (version 2.8.2). Peak calling was conducted using MACS2 software (version 2.1.1), and peak annotation and distribution analysis were performed using BEDtools (version 2.25.0). Differential peaks were identified using a Python script with Fisher’s exact test. To compare the differences in the common peaks shared between each group, we calculated the numbers of peaks in a 3-kb region (−2 kb to +1 kb around the transcription start site (TSS)) of each gene, as described in detail in the main text. The ATAC-seq data can be accessed in the Gene Expression Omnibus database under accession number GSE169502.

### RNA Preparation and Reverse Transcription-quantitative PCR (RT-qPCR)

Total RNA was extracted from SCC cells using TRIzol reagent (15596018; Invitrogen, CA, USA) according to the manufacturer’s instructions. Total RNA (1 µg) was used as a template for cDNA synthesis with a QuantScript RT Kit (KR103; Tiangen Biotech, Beijing, China). To determine mRNA expression levels, RT-qPCR was performed using a StepOnePlus Real-Time PCR system (Applied Biosystems, CA, USA) and PowerUp SYBR Green Master Mix (A25918; Applied Biosystems). LncRNA was reverse transcribed into cDNA using a lnRcute lncRNA First-Strand cDNA Kit (4992908; Tiangen Biotech), and lncRNA expression was quantified using a lnRcute lncRNA qPCR Kit (4992886; Tiangen Biotech). The relative expression of genes was calculated using the 2-^ΔΔCt^ method. The sequences of the primers used for RT-qPCR are listed in Table S2.

### Virus Production and Cell Infection

Full-length cDNAs were constructed using PCR amplification and subsequently inserted into the pLVX-IRES-neo vector (#632184; Clontech, CA, USA). The shRNA sequences were cloned into the pSIH1-puro vector (#26597; Addgene, MA, USA). Lentiviruses were produced in 293T cells with a second-generation packaging system containing psPAX2 (#12260; Addgene) and pMD2.G (#12259; Addgene). Transfection was performed using Hieff Trans Liposomal Transfection Reagent (40802; Yeasen, Shanghai, China). The shRNA and sgRNA sequences are listed in Table S3. Virus-containing medium was collected 48 and 72 h after transfection and was then added to cells supplemented with 8 mg/mL polybrene (TR-1003; Sigma) for lentiviral transduction. After 48 h, transduced cells were selected with 1 mg/mL puromycin (A610593; Sangon Biotech, Shanghai, China) or 200 mg/mL G418 (HY-17561; MCE) for 5-10 days and used for subsequent analysis.

### Cell Proliferation and Cell Viability Assays

Cell Counting Kit-8 reagent (CK04; Dojindo Laboratories, Kumamoto, Japan) was used to assess the cell proliferation rate or cell viability after DDP treatment as previously described (31).

### Cell Cycle Analysis

Cell cycle analysis was performed using a Cell Cycle Assay Kit (C543; Dojindo Laboratories) according to the manufacturer’s instructions. In brief, cells were harvested, washed with PBS, and fixed with 70% ice-cold ethanol for at least 2 h. Subsequently, the cells were washed and resuspended in 500 μL of assay buffer containing propidium iodide (PI; 1:20) and RNase (1:200) for 30 min at 37 °C in the dark and were then incubated for 30 min at 4 °C in the dark. Finally, the cell cycle distribution was analyzed using a flow cytometer.

### Caspase 3/7 Activity Assay

To assess DDP-induced apoptosis, caspase-3 and caspase-7 activities were measured using a Caspase Glo 3/7 Assay (G8090; Promega, WI, USA) according to the manufacturer’s instructions. In brief, SCC cells were seeded in 96-well plates and incubated with DDP at different concentrations for 24 h. Next, equal volumes of Caspase-Glo 3/7 reagent were added to the samples; after 1 h of incubation at room temperature, the luminescence of each sample was measured in a microplate luminometer. The blank reaction was used to measure background luminescence associated with the cell culture system and Caspase-Glo 3/7 reagent.

### Fluorescence In Situ Hybridization (FISH) and Immunofluorescence (IF)

To determine the localization of D-AS2 in SCC cells, FISH was performed using a Ribo^TM^ Fluorescent In Situ Hybridization Kit (C10910; RiboBio) according to the manufacturer’s instructions. In brief, cells were rinsed in PBS and fixed with 4% formaldehyde for 20 min at room temperature. Next, the cells were permeabilized in 0.1% Triton X-100 for 15 min at room temperature and washed twice with PBS. The cells were then incubated in prehybridization buffer for 30 min at 37 °C before hybridization. A probe targeting D-AS2 (labeled with Cy3 fluorescent dye) was used for hybridization in hybridization buffer for 12 h at 37 °C in a humidified chamber in the dark. After hybridization, the cells were sequentially washed at 42 °C in the dark with 4× SSC and 0.1% Tween 20 three times for 5 min each, with 2× SSC for 5 min, and with 1× SSC for 5 min. Finally, nuclei were stained with Hoechst 33342 for 15 min at room temperature in the dark, and images were acquired using laser scanning confocal microscopy. IF assays were performed according to previously described methods (30).

### Nuclear and Cytoplasmic Separation of RNA

Nuclear and cytosolic fractions were separated using a PARIS^TM^ kit (AM1921; Thermo Scientific) according to the manufacturer’s instructions. First, SCC cells were lysed with Cell Fraction Buffer to obtain the cytoplasmic components. Next, they were treated with Cell Disruption Buffer to obtain the nuclear components. Finally, RNA was extracted from the nuclear and cytoplasmic components. The expression levels of GAPDH, U6, and D-AS2 in the nucleus and cytoplasm of SCC cells were determined by RT-qPCR.

### RNA Pull-down Assay

D-AS2 sense and antisense strands were synthesized by in vitro transcription with a TranscriptAid T7 High Yield Transcription Kit (K0441; Thermo Scientific). The sequences were treated with RNase-free DNase I and purified using a GeneJET RNA Purification Kit (K0731; Thermo Scientific) according to the manufacturer’s instructions. Next, RNAs were labeled with a desthiobiotin tag by using a Pierce RNA 3′ End Desthiobiotinylation Kit (20163; Thermo Scientific). Pull-down of RNA– protein complexes was performed using a Pierce™ Magnetic RNA-Protein Pull-Down Kit (20164; Thermo Scientific). In brief, 50 pmol of labeled D-AS2 RNA and anti-D-AS2 RNA were bound to streptavidin magnetic beads, and 200 μg of protein was used for RNA binding. The RNA-protein bead mixture was incubated for 1 h at 4 °C with agitation. Finally, the RNA-binding proteins were eluted, and follow-up analysis was conducted.

### Mass Spectrometry (MS)

Eluted proteins were identified using a gel-based liquid chromatography–tandem mass spectrometry (Gel-LS-MS/MS) approach (Beijing Qinglian Biotech Co., Ltd., China). Mass spectra were analyzed using MaxQuant software (version 1.5.3.30) with the UniProtKB/Swiss-Prot human database.

### RIP Assay

The RIP assay was performed using a Magna RIP^TM^ RNA Binding Protein Immunoprecipitation Kit (17-700; Millipore) according to the manufacturer’s instructions. In brief, lysates of KYSE30 and NCI-H1703 cells were prepared in lysis buffer containing a protease inhibitor cocktail and an RNase inhibitor. Magnetic beads were preincubated with 5 μg of purified antibodies for 30 min at room temperature with rotation. Subsequently, the lysate supernatant was added to bead-labeled antibodies in immunoprecipitation buffer and incubated at 4 °C overnight. Finally, RNA was purified and subjected to relative quantitation using RT-qPCR. Normal rabbit IgG, as a negative control, was simultaneously assayed with antibodies against the target proteins. The IP efficiency was calculated using the percent input method. By using this method, we obtained signals from each immunoprecipitation reaction and expressed them as a percentage of the total input signal.

### CUT&RUN

CUT&RUN assays were performed using a CUT&RUN Assay Kit (86652; Cell Signaling Technology) according to the manufacturer’s instructions. In brief, cells were immobilized on concanavalin A magnetic beads to allow for subsequent buffer and reagent exchanges. Cell membranes were then permeabilized with digitonin to facilitate the nuclear entry of the primary antibody and pAG-MNase fusion enzyme. The target-specific primary antibody recruited pAG-MNase to chromatin through interactions between the antibody and the pAG domain of the fusion enzyme. The addition of Ca^2+^ activated pAG-MNase, which gently cleaved and liberated the desired chromatin fragments, allowing them to diffuse away from the genomic chromatin into the supernatant. Finally, DNA was purified using DNA purification spin columns (14209; Cell Signaling Technology) and was then identified and quantified using qPCR. The sequences of the primers used for the CUT&RUN assays are listed in Table S4.

### Detection of PLD Activity and PA Production

PLD activity and PA production were detected using a Phospholipase D Activity Assay Kit (Colorimetric) (183306, Abcam) and a Phosphatidic Acid Assay Kit (Fluorometric) (273335, Abcam), respectively, according to the manufacturer’s instructions. To prepare samples for PLD activity detection, 4 million adherent cells treated with or without inhibitors were collected and suspended in 100 μL of PLD buffer, incubated on ice for 30 min, and centrifuged at 14000 rpm for 3 min, and the supernatant was then collected for detection. To prepare samples for PA detection, 1 million adherent cells treated with or without inhibitors were collected and suspended in 1 mL of PA buffer, and lipids were then extracted with an organic solvent (chloroform/methanol/12 N HCl at a ratio of 2:4:0.1 v/v) for detection.

### Tag-lite Binding Assay

The saturation binding assays and competitive binding assays were performed according to the manufacturer’s instructions for the PerkinElmer Tag-lite binding assay. The SNAP-tagged FPR1 or FPR2 plasmids were constructed from Tag-lite pT8-SNAP (Cisbio Bioassays; Codolet, France), pEZ-Lv105-FPR1 (#Z7406; GeneCopoeia, GuangZhou, China) and pEZ-Lv105-FPR2 (#A0060; GeneCopoeia). In brief, 293T cells or SCC cells were transfected with SNAP-tagged FPR1 or FPR2 overnight at 37 °C in 5% CO_2_. Cells expressing each SNAP-tag-GPCR protein were labeled with 100 nM Tag-lite SNAP-Lumi4-Tb for 1 h at 37 °C in 5% CO_2_.

A saturation binding assay was performed to measure the total and nonspecific binding of increasing concentrations of XL665-tagged FAM3D under equilibrium conditions. Labeled FAM3D was titrated into a solution containing 20 K labeled cells per well and incubated to equilibrium. The HTRF ratio determined from this titration represents the total binding. A negative control reaction using unlabeled ligand (FPR Agonist 43) was included to account for nonspecific binding of the labeled ligand to the receptor, nonspecific molecules, and the microplate. In this reaction, labeled FAM3D was titrated into a solution containing 20 K labeled cells and a 100-fold molar excess of (10 µM) FPR Agonist 43. The HTRF ratio determined from this titration represents the nonspecific binding. The specific binding was determined by subtracting the nonspecific binding from the total binding at each labeled FAM3D concentration. A competitive binding assay was performed to measure the dissociation constant, Ki. The competitors FPR Agonist 43 and HCH6-1 for FPR1 were titrated into a solution containing 8 nM (80% Kd) labeled FAM3D and 20 K labeled cells; the competitors FPR Agonist 43 and PBP10 TFA for FPR2 were titrated into a solution containing 16 nM (80% Kd) labeled FAM3D and 20-K labeled cells. The ratio of the acceptor and donor emission signals for each individual well was calculated as follows: Ratio = (Signal 665 nm)/(Signal 615 nm) × 10^4. The binding curves were fitted using GraphPad Prism 8.

## Statistical Analysis

Two-tailed Student’s *t* test, two-way ANOVA, or the Kaplan–Meier method was used for statistical analyses. For all statistical analyses, differences with *P* < 0.05 were considered statistically significant, and data from at least three biologically independent experiments with similar results are reported. The sample size for each statistical analysis was determined based on pretests and previous similar experiments. Data analyses were performed using GraphPad Prism (version 8.01; GraphPad Software, CA, USA).

## Results

### Identification of D-AS2 as a critical lipid-related lncRNA in SCC cell chemoresistance

Long noncoding RNAs (lncRNAs) have been shown to participate in nearly all kinds of biological processes—both physiological processes, such as differentiation and development (32, 33), and processes related to diseases such as cancer and cardiac disease (34, 35). However, the critical lncRNAs that mediate specific functions in each context remain largely unexplored. To define the mechanisms underlying cancer cell chemoresistance, we treated SCC cells with several rounds of high-dose cisplatin (DDP) to mimic the clinical treatment regimen used for cancer patients, thus establishing two clinical mimic DDP-resistant cell models: YES2/DDP (31) and KYSE450/DDP (Fig. 1A). As expected, cell viability assays indicated that YES2/DDP and KYSE450/DDP cells showed stronger resistance to DDP treatment than the corresponding parental cells (Fig. S1A). In addition, the chemoresistant cells showed significantly lower proliferation rates than the parental cells (Fig. S1B), consistent with the characteristics of some previously reported chemoresistant cancer cells (36–38).

**Figure 1.**
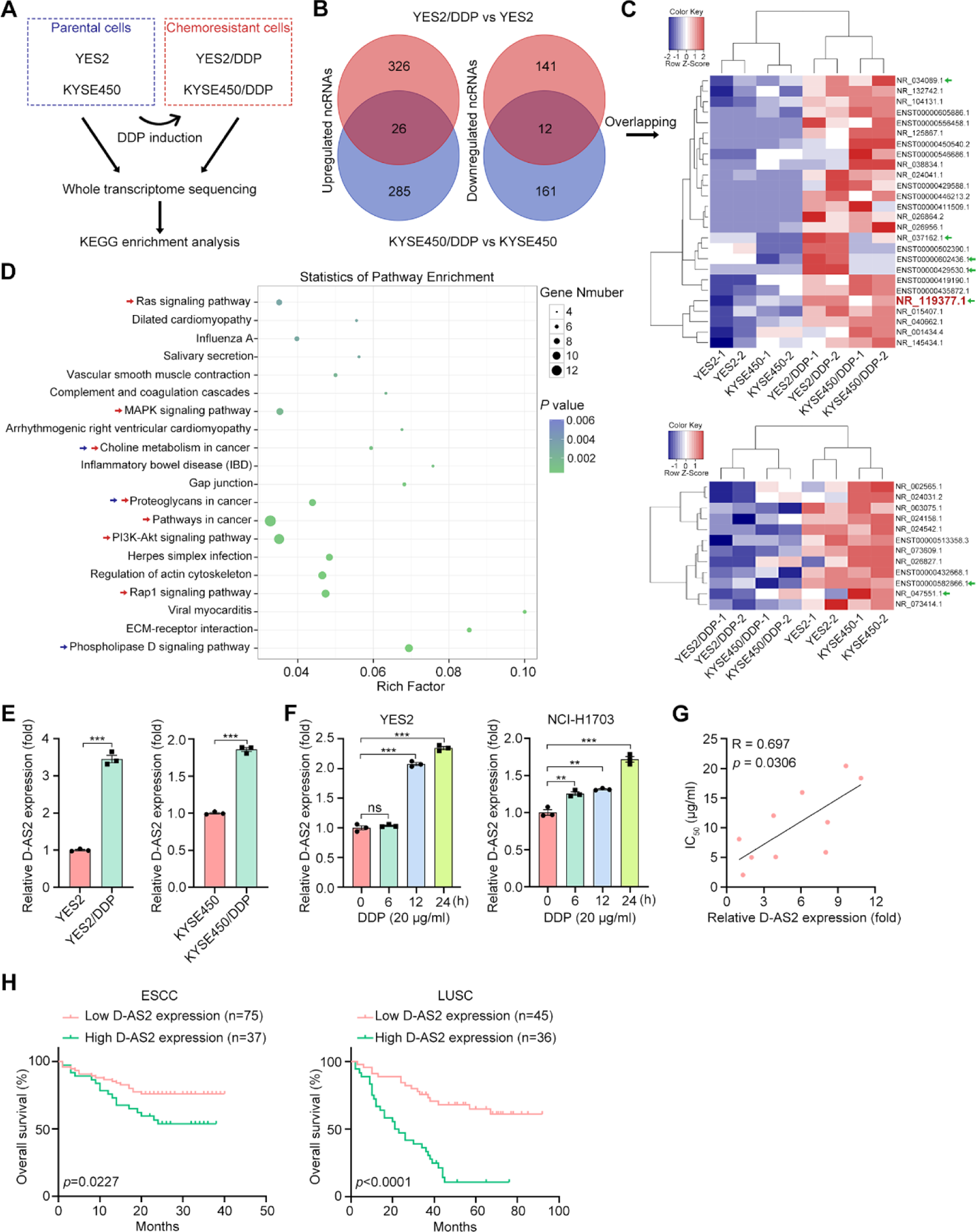
D-AS2 is elevated in chemoresistant SCC cells and is associated with metabolic events. (**A**) Schematics showing the strategy for deciphering the molecular mechanisms underlying chemoresistance in SCC. (**B**) Numbers of significantly upregulated and downregulated ncRNAs in the two chemoresistant cell lines compared to the corresponding parental lines. (**C**) Heatmap showing the expression profile of the 26 ncRNAs upregulated and 12 ncRNAs downregulated in both chemoresistant cell lines. The green arrows indicate ncRNAs that were enriched in PLD signaling, as determined by their predicted target genes. (**D**) Top 20 significantly enriched pathways identified by KEGG analysis of the consistently differentially expressed mRNAs in both chemoresistant cell models. The red arrows indicate cancer-related pathways, and the blue arrows indicate metabolic pathways. (**E**) RT-qPCR detection of D-AS2 expression in the parental and chemoresistant cell models. The data are presented as the mean ± s.d. values; two-tailed *t* test, \*\*\**P* < 0.001; n = 3. (**F**) RT-qPCR detection of D-AS2 expression in YES2 and NCI-H1703 cells treated with DDP for 0, 6, 12, and 24 h. The data are presented as the mean ± s.d. values; two-tailed *t* test, \*\*\**P* < 0.001, \*\**P* < 0.01, ns: not significant; n = 3. (**G**) Correlation analysis between the DDP IC_50_ and D-AS2 expression in SCC cell lines. Spearman correlation coefficients are shown. (**H**) Overall survival of esophageal and lung SCC patients with low (below mean) or high (above mean) D-AS2 expression. Kaplan– Meier survival plots are shown.

To further assess the characteristics of the chemoresistant SCC cells and identify crucial genes that contribute to chemoresistance, we performed whole-transcriptome sequencing of YES2/DDP, KYSE450/DDP, YES2, and KYSE450 cells and analyzed the differentially expressed mRNAs and ncRNAs in the chemoresistant and parental cell lines (Fig. 1A). The numbers of upregulated mRNAs and ncRNAs were markedly greater than the numbers of downregulated mRNAs and ncRNAs in the chemoresistant cells in both pairs of cell lines, indicating that more genes were activated than suppressed in the chemoresistant cells (Fig. S1C). In total, 352/153 and 311/173 significantly upregulated/downregulated ncRNAs were identified in YES2/DDP and KYSE450/DDP cells, respectively, and 26/12 of these significantly upregulated/downregulated ncRNAs showed consistent upregulation/downregulation in both chemoresistant cell lines (Figs. 1B and 1C). KEGG enrichment analysis of the consistently differentially expressed mRNAs in the two pairs of chemoresistant cell lines indicated enrichment of these mRNAs in several cancer-regulated pathways (Fig. 1D), consistent with the significant changes in the chemoresistant and proliferative phenotypes observed in our chemoresistant models. Interestingly, the differentially expressed mRNAs were also enriched in a few metabolic pathways, and PLD signaling was the most significantly enriched pathway (Fig. 1D). Moreover, KEGG analysis of the predicted targets of the 38 consistently differentially regulated ncRNAs identified enrichment in a panel of metabolism-related pathways and the platinum drug resistance pathway (Fig. S1D). Again, PLD signaling was the most significantly enriched specific pathway (Fig. S1D). These results strongly suggest that metabolic reprogramming occurs during the acquisition of chemoresistance.

Next, we identified the ncRNA that plays an essential role in chemoresistance and metabolic reprogramming. First, we analyzed the involvement of the 38 consistently regulated ncRNAs in PLD signaling. A total of 7 ncRNAs (5 upregulated, 2 downregulated; indicated by the green arrows in Fig. 1C) were enriched in PLD signaling, as indicated by their predicted target genes. Activated ncRNAs in chemoresistant cells are potentially effective therapeutic targets; thus, we focused mainly on the upregulated ncRNAs for subsequent validation. Since the functional effects of ncRNAs usually rely on an adequate expression level, we then evaluated the abundance of the 5 upregulated ncRNAs and found that only NR_119377.1 (i.e., D-AS2) showed an adequate abundance in the tested cells (reads per kilobase transcript per million mapped reads >1 in parental cells and >10 in chemoresistant cells). The elevation of D-AS2 expression was confirmed using RT–qPCR analysis in another independent culture of chemoresistant and parental cells (Fig. 1E). To validate whether upregulation of D-AS2 is an essential event that contributes to chemoresistance instead of an event accompanying acquired chemoresistance after long-term culture, we treated YES2 and KYSE450 cells with DDP for a short duration (6, 12, or 24 h) and evaluated the expression of D-AS2. D-AS2 expression was elevated in a time-dependent manner after DDP exposure (Fig. 1F), indicating that elevation of D-AS2 expression plays an essential role in both short- and long-term exposure to cytotoxic drugs. To extend the application of our results, we determined the D-AS2 expression level (Fig. S1E) and half-maximal inhibitory concentration (IC_50_) of DDP (Fig. S1F) in several SCC cell lines derived from SCCs of the skin, tongue, esophagus, and lung. Correlation analysis indicated a significant positive correlation between the IC_50_ values and D-AS2 expression levels in the cell lines (Fig. 1G). Finally, we performed ISH to detect D-AS2 expression in two cohorts of esophageal and lung SCCs. D-AS2 expression was significantly correlated with poor prognosis in both SCC cohorts (Fig. 1H) but did not differ between normal tissue and SCC samples (Fig. S1G).

### D-AS2 overexpression phenocopies the chemoresistant characteristics of chemoresistant cell models

Next, we performed functional studies to elucidate the role of D-AS2 in the chemoresistance of SCC cells. First, we designed three independent ASO sequences and validated their similar efficiency in silencing D-AS2 expression (Fig. S2A). Subsequently, we transfected the two chemoresistant SCC cell lines with D-AS2-targeting or negative control ASOs to evaluate their sensitivity to DDP treatment. The cell viability assays showed that silencing D-AS2 sensitized both chemoresistant cell lines to DDP treatment (Fig. 2A). We also silenced D-AS2 in two other parental SCC cell lines, NCI-H1703 and KYSE30, which have relatively high D-AS2 expression levels and DDP IC_50_ values (Figs. S1E and S1F). Knockdown of D-AS2 reduced the resistance of NCI-H1703 and KYSE30 cells to DDP (Fig. 2B). Conversely, exogenous expression of D-AS2 in the three SCC cell lines with relatively low D-AS2 expression significantly increased their resistance to DDP-induced cell death (Figs. S2B and S2C). Surprisingly, in contrast to the observations for most RNAs or proteins that showed consistent oncogenic or tumor-suppressive effects in a specific cancer type, D-AS2 overexpression significantly reduced the proliferation of SCC cells (Fig. S2D). However, this phenotype suggested that D-AS2 overexpression completely phenocopied the characteristics of our established chemoresistant cell models.

**Figure 2.**
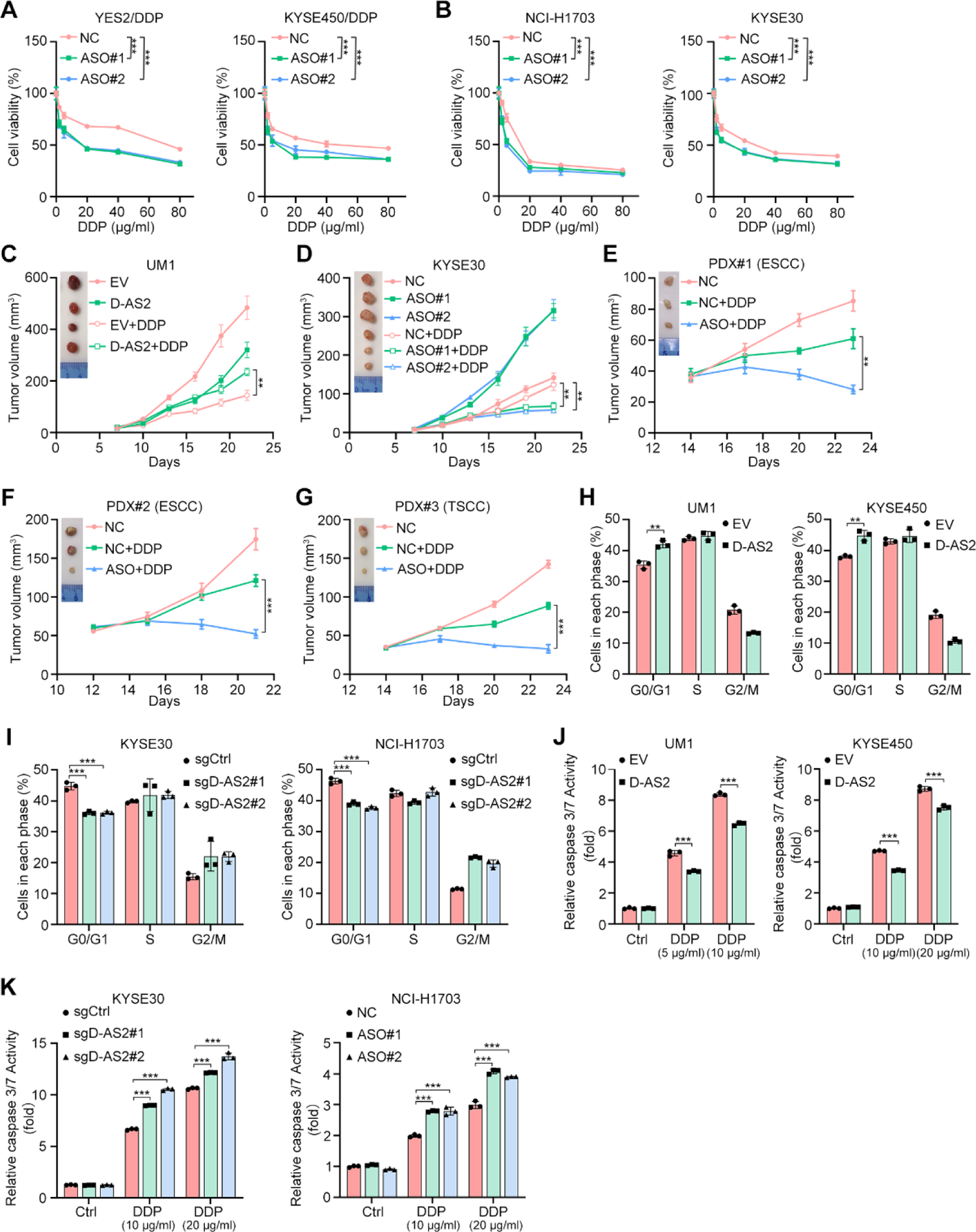
D-AS2 induces cell cycle arrest and chemoresistance in SCCs. (**A, B**) Relative viability of the indicated cell lines transfected with negative control (NC) or D-AS2 (ASO#1, ASO#2) ASOs after DDP treatment for 24 h. The data are presented as the mean ± s.d. values; two-way ANOVA, \*\*\**P* < 0.001; n = 5. (**C**) Representative images and tumor volumes of xenografts derived from UM1 cells expressing empty vector (EV) or D-AS2. The data are presented as the mean ± s.e.m. values; two-tailed *t* test, \*\**P* < 0.01; n = 8. (**D**) Representative images and volumes of xenografts derived from KYSE30 cells and treated with NC or D-AS2 (ASO#1, ASO#2) ASOs in vivo. The data are presented as the mean ± s.e.m. values; two-tailed *t* test, \*\**P* < 0.01; n = 8. (**E-G**) Representative images and volumes of xenografts in in vivo PDX models treated with NC or D-AS2 (ASO) ASOs. The data are presented as the mean ± s.e.m. values; two-tailed *t* test, \*\*\**P* < 0.001, \*\**P* < 0.01; n = 8 (except for n = 7 in the NC + DDP arm and n = 4 in the ASO + DDP arm of PDX#1). (**H, I**) Cell cycle analyses of the indicated cells. The data are presented as the mean ± s.d. values; two-tailed *t* test, \*\*\**P* < 0.001, \*\**P* < 0.01; n = 3. (**J, K**) Detection of caspase 3/7 activity in the indicated cells. The data are presented as the mean ± s.d. values; two-tailed *t* test, \*\*\**P* < 0.001; n = 3.

To elucidate the functions of D-AS2 in vivo, we subcutaneously xenografted control or D-AS2-overexpressing UM1 cells into nude mice and administered saline or DDP one week later. Consistent with the in vitro experimental results, D-AS2-overexpressing UM1 cells showed a reduced growth rate and exhibited increased resistance to DDP treatment (Fig. 2C). Conversely, in vivo application of ASOs targeting D-AS2 in KYSE30 xenografts promoted the growth of these xenografts and sensitized them to DDP (Fig. 2D). To exclude the potential off-target effects of ASO application in vivo, we designed two independent sgRNA sequences targeting D-AS2, which led to moderate and significant depletion of D-AS2 expression in SCC cells when introduced via lentiviral transduction (Figs. S2E and S2F). Next, we repeated the xenograft assays with KYSE30 and NCI-H1703 cells that stably expressed control or D-AS2-targeting sgRNAs. The phenotypes of tumor growth and chemoresistance were affected in a dose-dependent manner by D-AS2 depletion (Figs. S2G and S2H), validating the on-target effect of D-AS2 targeting. To further determine the effect of interfering with D-AS2 during SCC chemotherapy, we established three PDX models from tissue samples of tongue and esophageal SCC and administered ASOs in vivo to mimic clinical treatment. Strikingly, in all three models, in vivo administration of D-AS2-targeting ASOs significantly sensitized SCC samples to DDP treatment (Figs. 2E-G). These results provide a promising approach for D-AS2 interference by using ASOs or other feasible methods to promote therapeutic effects of cytotoxic chemotherapeutic agents.

The proliferation rate of cells depends mainly on cell cycle progression; thus, we determined whether D-AS2 affects the cell cycle in SCC cells. To this end, we first evaluated the cell cycle distribution of control and D-AS2-overexpressing UM1 and KYSE450 cells. Notably, more D-AS2-overexpressing cells than control cells accumulated in G0/G1 phase (Figs. 2H and S2I). On the other hand, D-AS2 depletion led to a reduction in the G0/G1-phase population and an increase in the G2/M-phase population in both KYSE30 and NCI-H1703 cells (Figs. 2I and S2J). Next, we assessed caspase activation to determine the effect of DDP on cell death. As expected, D-AS2 overexpression suppressed the activation of caspase 3/7 after DDP treatment (Fig. 2J), and both sgRNA- and ASO-mediated D-AS2 silencing led to increased caspase activity in DDP-treated SCC cells (Fig. 2K).

### FAM3D acts as a lipid-related target to mediate the function of D-AS2

To elucidate the downstream events underlying the functions of D-AS2, we performed RNA-seq in control and D-AS2-overexpressing KYSE450 cells. Surprisingly, unlike the findings in chemoresistant cells, in which more genes were upregulated than downregulated, D-AS2 overexpression led to significant suppression of the transcriptome in SCC cells (Fig. 3A). These results indicate that D-AS2 contributes mainly to the downregulation of the downstream targets in chemoresistant cells. The gene set enrichment analysis (GSEA) results indicated a negative correlation between apoptosis and D-AS2 expression (Fig. S3A), consistent with our previous speculation that D-AS2 overexpression suppresses cell death after DDP treatment. Interestingly, although both glycolysis and hypoxia were negatively correlated with D-AS2 overexpression, no significant correlation was found between D-AS2 expression and OXPHOS (Fig. S3B). The enrichment of several metabolic pathways also suggested the critical role of D-AS2 in the metabolic reprogramming of SCC cells (Fig. S3C).

**Figure 3.**
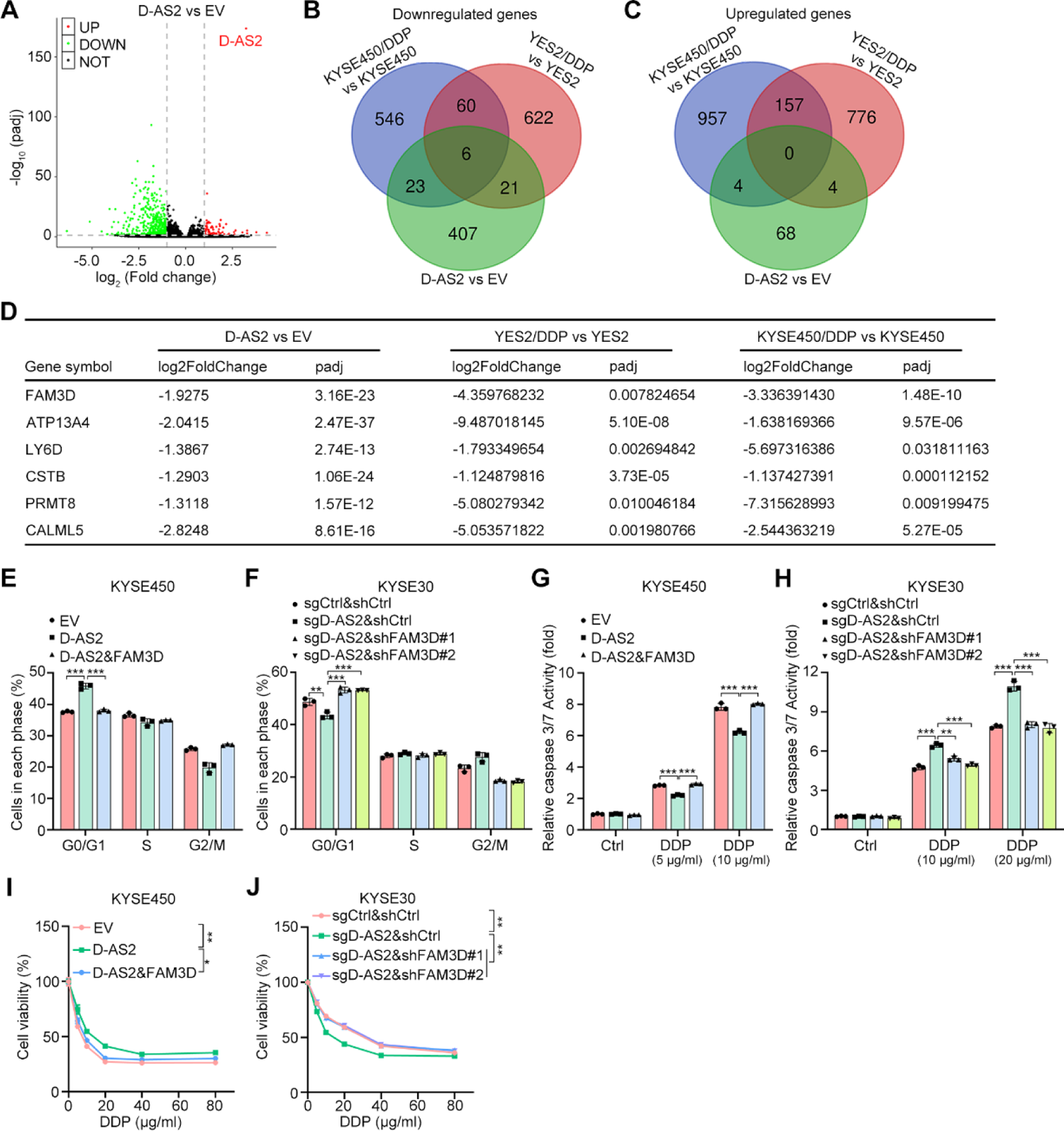
FAM3D acts as a lipid-related target to mediate the function of D-AS2. (**A**) Volcano plots showing upregulated and downregulated mRNAs in D-AS2-overexpressing KYSE450 cells compared to control cells. (**B, C**) Number of genes that showed significant downregulation (B) and upregulation (C) in D-AS2-overexpressing and chemoresistant cell models. (**D**) Fold changes in gene expression in D-AS2-overexpressing and chemoresistant cell models. (**E, F**) Cell cycle analyses of the indicated cells. The data are presented as the mean ± s.d. values; two-tailed *t* test, \*\*\**P* < 0.001, \*\**P* < 0.01; n = 3. (**G, H**) Detection of caspase 3/7 activity in the indicated cells. The data are presented as the mean ± s.d. values; two-tailed *t* test, \*\*\**P* < 0.001, \*\**P* < 0.01; n = 3. (**I, J**) Relative viability of the indicated cell lines after DDP treatment for 24 h. The data are presented as the mean ± s.d. values; two-way ANOVA, \*\**P* < 0.01, \**P* < 0.05; n = 3.

We next identified the critical downstream target that is directly regulated by D-AS2 and contributes to chemoresistance in SCC. First, we analyzed the genes that were consistently upregulated or downregulated in D-AS2-overexpressing cells and both chemoresistant cell models (Fig. 3B and 3C). Six genes were consistently downregulated in all the three comparison groups (Fig. 3B and 3D), while no consistently upregulated gene was identified (Fig. 3C). Among the six downregulated genes, we focused on *FAM3D* because the encoded extracellular secreted protein activates intracellular signaling as a ligand of cell membrane receptors such as GPCRs (39). Accordingly, the ELISA results indicated that D-AS2 suppressed extracellular FAM3D protein secretion (Fig. S3D).

We then determined whether FAM3D is critical for the functions of D-AS2 in the cell cycle and chemoresistance. Cell cycle analysis showed that exogenous expression of FAM3D abolished the effect of D-AS2 on G0/G1 arrest in KYSE450 cells (Figs. 3E, S3E, and S3G). Consistently, silencing FAM3D by using two independent shRNA sequences rescued the effect of D-AS2 depletion on cell cycle progression (Figs. 3F, S3F, and S3H). FAM3D overexpression increased the activation of caspase 3/7 in D-AS2-overexpressing KYSE450 cells (Fig. 3G), whereas FAM3D knockdown suppressed DDP-induced activation of caspase 3/7 in KYSE30 cells with D-AS2 depletion (Fig. 3H). Similarly, cell viability assays showed that FAM3D rescued the effects of D-AS2 on the chemosensitivity of SCC cells to DDP (Figs. 3I and 3J). Taken together, these results indicate that FAM3D is the direct downstream target of D-AS2 and contributes to the effects of D-AS2 on cell cycle progression and chemoresistance to cytotoxic drugs.

### D-AS2 associates with histones to regulate the distal element of FAM3D

To explore the mechanism by which D-AS2 regulates FAM3D and, subsequently, metabolic reprogramming and chemoresistance, we assessed the subcellular distribution of D-AS2 in SCC cells. FISH assays showed that D-AS2 was localized mainly in the nucleus of SCC cells (Fig. 4A). Nuclear-cytoplasmic fractionation followed by RT-qPCR analysis also confirmed the dominant nuclear subcellular localization of D-AS2 (Fig. S4A). These results indicated that D-AS2 might function by directly affecting transcriptional regulation events. Next, we performed RNA pull-down assays followed by MS analysis. To exclude false positive results of MS analysis, we repeated the experiments twice independently in KYSE30 cells and performed a separate experiment in NCI-H1703 cells. The MS analysis results suggested that several histones were associated with D-AS2. However, no specific histone was detected in any of the three independent MS analyses (Fig. 4B). More surprisingly, both core and linker histones were identified (Fig. 4B). To validate our MS results, we performed RIP assays by using antibodies against the linker histones H1.3 and H1.5 and the core histones H2A, H2B, H3, and H4. D-AS2 interacted with all linker and core histones detected in both KYSE30 and NCI-H1703 cells (Fig. 4C). To further evaluate the specificity of our RNA pull-down assays, we repeated these experiments with both the sense and antisense sequences of D-AS2 and detected histones by immunoblot (IB) analysis using specific antibodies. Linker and core histones were specifically associated with D-AS2 but not its antisense sequence (Fig. 4D). These results indicate that D-AS2 generally and dynamically interacts with histones, which explains the MS results that no specific histone subtype was identified. Modifications of histones, such as acetylation and methylation, have been shown to play essential roles in the regulation of gene transcription and are thus called histone marks (40, 41). However, despite the associations between histones and D-AS2, the levels of histone marks remained unchanged following either depletion or overexpression of D-AS2 (Fig. S4B). Thus, we speculated that D-AS2 might regulate gene transcription by regulating the chromatin structure instead of by affecting the abundance of histone marks. To confirm this hypothesis, we performed ATAC-seq in control and D-AS2-silenced NCI-H1703 cells. The total ATAC-seq signal around the TSS region of all genes did not change markedly after D-AS2 knockdown (Fig. S4C), supporting our hypothesis that D-AS2 mediates gene transcription by modulating distal elements instead of promoter regions. Next, we analyzed the total numbers of ATAC-seq peaks in the different groups to assess the effects of D-AS2 on the whole chromatin structure in SCC cells. Strikingly, silencing of D-AS2 caused the loss of specific peaks in 2447 genes and the gain of specific peaks in 1753 genes, suggesting a significant effect of D-AS2 on the transcriptome (Fig. 4E). However, in contrast to the ubiquitous suppression of mRNA expression after D-AS2 overexpression, fewer unique peaks were noted in D-AS2-knockdown cells than in control cells (Fig. 4E). These results contradicted our RNA-seq data, which indicated that D-AS2 triggers global suppression of the transcriptome. To address the controversial results of ATAC-seq and RNA-seq indicating an activated or suppressed transcriptional status, respectively, we focused on the 20876 genes with common peaks shared by the different groups in the ATAC-seq results (Fig. 4E). To determine the differences in the genes with shared peaks, we first set the length of all genes to 3 kb (−2 kb to +1 kb around the TSS) and determined the numbers of peaks in this 3-kb region of each gene. Next, we compared the numbers of peaks in these genes (within the 3-kb region) and identified the differentially expressed genes. By using this method, we found more upregulated genes than downregulated genes in D-AS2-knockdown cells (Fig. 4F), consistent with the RNA-seq data.

**Figure 4.**
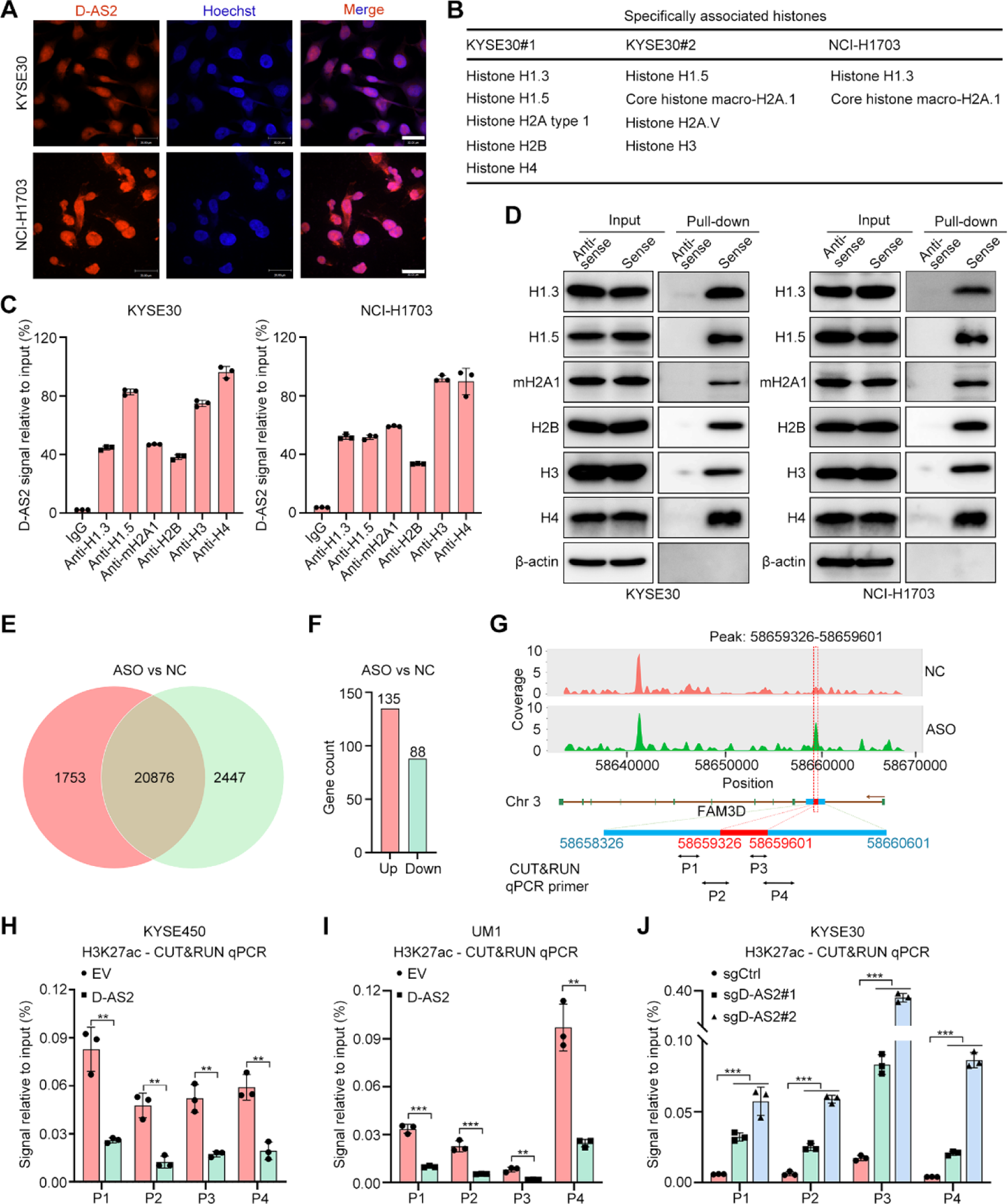
D-AS2 reduces H3K27ac enrichment at the FAM3D enhancer region. (**A**) FISH detection of D-AS2 localization in KYSE30 and NCI-H1703 cells. Scale bar, 30 μm. (**B**) Lists of D-AS2-interacting histones identified using MS in KYSE30 and NCI-H1703 cells. (**C**) RT-qPCR measurement of the D-AS2 level in precipitates from RIP assays performed using IgG or specific antibodies. The data are presented as the mean ± s.d. values; n = 3. (**D**) IB detection of histones in precipitates from RNA pull-down assays performed using antisense or sense sequences of D-AS2. (**E**) Numbers of genes that exhibited shared or unique peaks in D-AS2-silenced and control NCI-H1703 cells. (**F**) Numbers of genes with significant differences in the number of peaks in the gene collections that exhibited shared peaks. (**G**) Unique peak identified in D-AS2-silenced NCI-H1703 cells. (**H-J**) CUT&RUN assay of the FAM3D enhancer region in D-AS2-depleted and D-AS2-overexpressing SCC cells with antibodies against H3K27ac. The data are presented as the mean ± s.d. values; two-tailed *t* test, \*\*\**P* < 0.001, \*\**P* < 0.01; n = 3.

To further determine the mechanism by which D-AS2 regulates FAM3D expression, we analyzed ATAC-seq signals at the genomic locus of FAM3D. In D-AS2-knockdown cells, a unique H3K27ac peak was detected around an enhancer region in FAM3D (42) (Fig. 4G), indicating that FAM3D might be activated by increased accessibility to the transcriptional activation mark H3K27ac. Thus, we performed CUT&RUN assays to determine the effects of D-AS2 on the association between the FAM3D enhancer region and H3K27ac and found that D-AS2 overexpression decreased H3K27ac enrichment in the FAM3D enhancer region in KYSE450 and UM1 cells (Figs. 4H and 4I). On the other hand, D-AS2 depletion in KYSE30 cells increased H3K27ac enrichment in the FAM3D enhancer region (Fig. 4J). In contrast, another histone mark that usually correlates with enhancers, H3K4me1, showed no consistent change upon D-AS2 overexpression or depletion (Figs. S4D-F). Taken together, these results show that D-AS2 hinders FAM3D transcription by reducing H3K27ac in the enhancer region.

### D-AS2 enhances PLD activity via FAM3D-mediated FPR1 and FPR2 signaling

We next determined whether D-AS2/FAM3D regulates PLD activity and the subsequent production of PA. Indeed, overexpression of D-AS2 markedly increased both PLD activity and PA production, and exogenous expression of FAM3D blocked the enhancing effects of D-AS2 on PLD activity and PA production (Figs. 5A and 5B). The D-AS2 depletion-mediated reductions in PLD activity and PA production were also reversed by shRNA-mediated FAM3D silencing (Figs. 5A and 5B).

**Figure 5.**
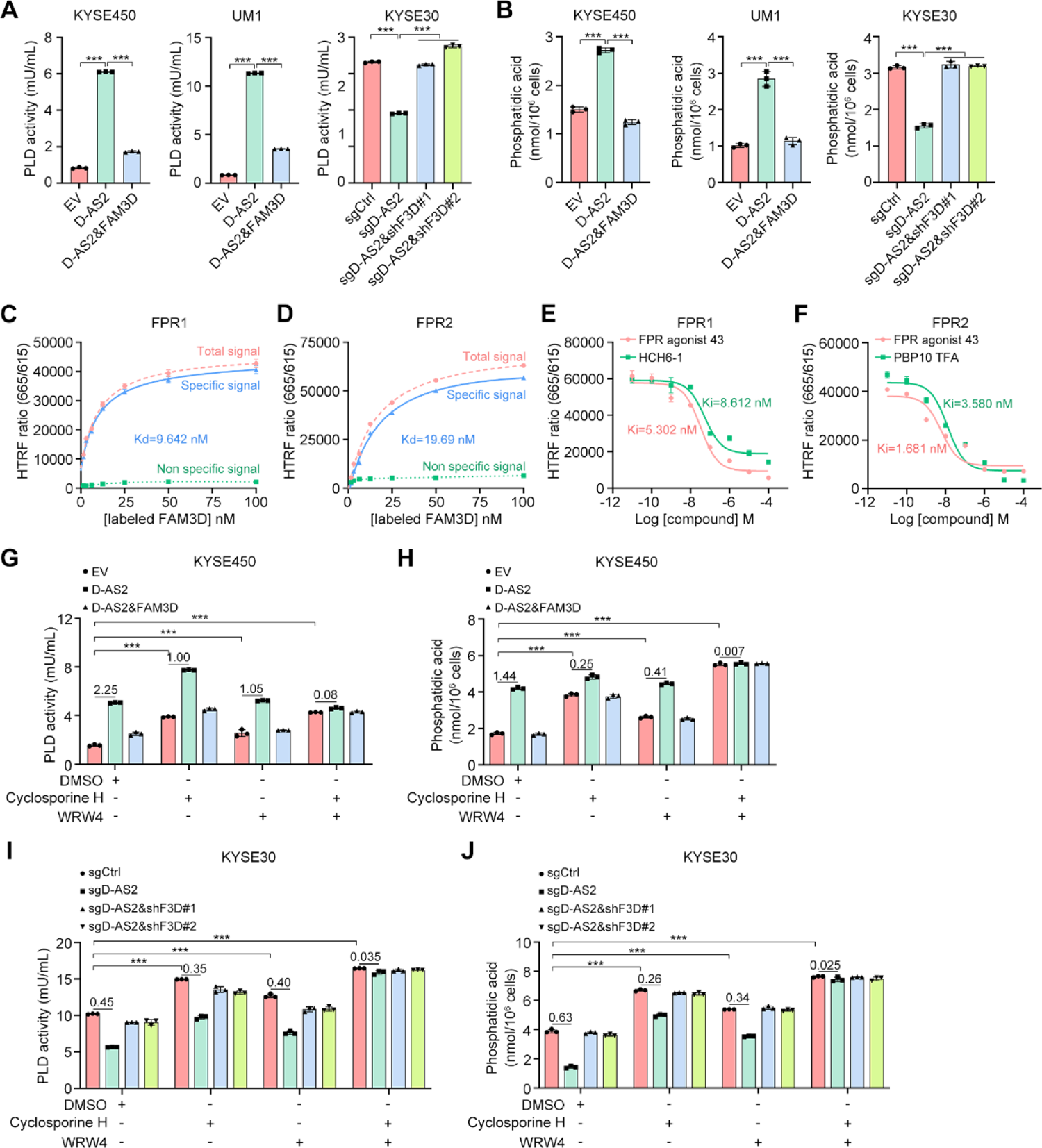
FAM3D associates with FPR1 and FPR2 to suppress PLD signaling. (**A**) Detection of PLD activity in the indicated cells. The data are presented as the mean ± s.d. values; two-tailed *t* test, \*\*\**P* < 0.001, \*\**P* < 0.01; n = 3. (**B**) Detection of PA production in the indicated cells. The data are presented as the mean ± s.d. values; two-tailed *t* test, \*\*\**P* < 0.001; n = 3. (**C, D**) Saturation curve of FAM3D binding to FPR1 and FPR2. 293T cells transiently expressing FPR1- and FPR2-tagged SNAP were incubated with increasing concentrations of labeled FAM3D. A prominent homogeneous time-resolved fluorescence (HTRF) signal is visible. Nonspecific binding was measured by adding 10 µM FPR Agonist 43 to the wells. The dissociation constants (K_d_ values) are shown. The data are presented as the mean ± s.d. values; n = 3. (**E, F**) Competitive binding curve for FAM3D to FPR1 and FPR2. 293T cells transiently expressing FPR1- and FPR2-tagged SNAP were incubated with increasing concentrations of the indicated competitors. The concentration of labeled FAM3D used for FPR1 was 8 nM and that used for FPR2 was 16 nM. The inhibitory constants (K_i_ values) are shown. The data are presented as the mean ± s.d. values; n = 3. (**G-J**) Detection of PLD activity and PA production in the indicated cells. Cells were treated with cyclosporine H (5 μM; FPR1 inhibitor) or WRW4 (10 μM; FPR2 inhibitor) for 1 h. The fold increases after D-AS2 overexpression and fold decreases after D-AS2 depletion are shown. The data are presented as the mean ± s.d. values; two-tailed *t* test, \*\*\**P* < 0.001; n = 3.

Next, we sought to investigate the underlying mechanism by which FAM3D mediates PLD activation by D-AS2. Previous reports indicate that a wide variety of hormones, growth factors, and cytokines activate PLD via stimulation of G protein-coupled receptors (GPCRs) and receptor tyrosine kinases (RTKs) (43). One study revealed FAM3D as a new endogenous chemotactic agonist for formyl peptide receptor (FPR) 1 and FPR2, both of which are Gα_i_ protein coupled seven-transmembrane-domain GPCRs (39). Based on the structural and functional properties of their α-subunits, heterotrimeric G proteins are subclassified into 4 major families: G_s_, G_i/o_, G_q/11_, and G_12/13_ (44). The Gαi coupled GPCRs negatively regulate key effectors and the generation of secondary messengers (45, 46). First, to confirm that FAM3D acts as a ligand that can specifically bind FPR1 and FPR2, we performed tag-lite binding assays in 293T cells. The results of the saturation binding assays showed that upon the addition of increasing concentrations of labeled FAM3D, both the FPR1-FAM3D and FPR2-FAM3D binding signals were increased in a dose-dependent manner (Figs. 5C and 5D). Specifically, the dissociation constants were calculated to be 9.642 nM for FPR1 and 19.69 nM for FPR2, indicating the higher affinity of FAM3D for FPR1 (Figs. 5C and 5D). Additionally, competitive binding assays were performed with two competitors for each receptor: FPR Agonist 43 and HCH6-1 for FPR1 and FPR Agonist 43 and PBP10 TFA for FPR2. Upon the addition of increasing concentrations of the competitor compounds, the binding signals between labeled FAM3D and the GPCRs decreased gradually with increasing competitor concentration (Figs. 5E and 5F). These results were further confirmed in SCC cell lines KYSE30 and KYSE450 (Figs. S5A-D).

Finally, we determined whether D-AS2/FAM3D-regulated PLD activity is mediated by FPR1 and FPR2 signaling. KYSE450 cells with ectopic D-AS2 or D-AS2 and FAM3D were treated with cyclosporine H (a specific inhibitor of FPR1) or WRW4 (a specific inhibitor of FPR2). The overexpression of D-AS2 dramatically increased PLD activity (by 2.25-fold) and PA production (by 1.44-fold), and ectopic expression of FAM3D almost completely abolished these promotive effects (Figs. 5G and 5H). Either cyclosporine H or WRW4 treatment alone partially reduced the promotive effects of D-AS2 on PLD activity and PA production (Figs. 5G and 5H). When FPR1 and FPR2 were simultaneously inhibited, the PLD signaling was strikingly activated but not further promoted by D-AS2 (Figs. 5G and 5H). Consistent with this finding, in KYSE30 cells expressing the D-AS2-targeting sgRNA or FAM3D-targeting shRNAs, the fold decreases in PLD activity and PA production mediated by D-AS2 depletion were reduced after inhibition of either FPR1 or FPR2 (Figs. 5I and 5J). Furthermore, synergistic inhibition of FPR1 and FPR2 almost completely abolished the suppressive effects of D-AS2 depletion on PLD activity and PA production (Figs. 5I and 5J). Taken together, these results indicate that both FPR1 and FPR2 play important roles in D-AS2/FAM3D-mediated downstream signaling.

### D-AS2 promotes PLD/PA-mediated YAP activation

Since PA has been reported to regulate YAP signaling (22, 23), we next determined the role of D-AS2 on YAP activation. Overexpression of D-AS2 in KYSE450 cells dramatically suppressed YAP phosphorylation at S127 and decreased LATS1 phosphorylation at S909 (Fig. S6A). D-AS2 also promoted the transcription of the YAP downstream genes CTGF and CYR61, suggesting a broad role of D-AS2 in Hippo pathway regulation (Fig. S6B). Overexpression of D-AS2 in UM1 cells obtained consistent results (Figs. S6C and S6D). On the other hand, silencing of D-AS2 in KYSE30 cells increased the phosphorylation of YAP and LATS1 (Fig. S6E). To further determine whether PLD is required for D-AS2-mediated YAP activation, we treated control and D-AS2-overexpressing KYSE450 and UM1 cells with the PLD inhibitors FIPI and CAY10594. Treatment with PLD inhibitors markedly increased YAP and LATS1 phosphorylation in both groups and completely abolished the suppressive effect of D-AS2 on YAP activity (Figs. 6A and S6F). Consistent with these findings, inhibitor treatment completely suppressed the transcriptional activation of YAP targets by D-AS2 (Figs. 6B and S6G). YAP phosphorylated at S127 by activated LATS1/2 is sequestered in the cytoplasm, and unphosphorylated YAP translocates into the nucleus to activate gene transcription (22). Thus, we next analyzed the subcellular localization of YAP in KYSE450 cells and found that D-AS2 markedly induced YAP nuclear translocation, while treatment with FIPI and CAY10594 abolished this effect (Figs. 6C and 6D).

**Figure 6.**
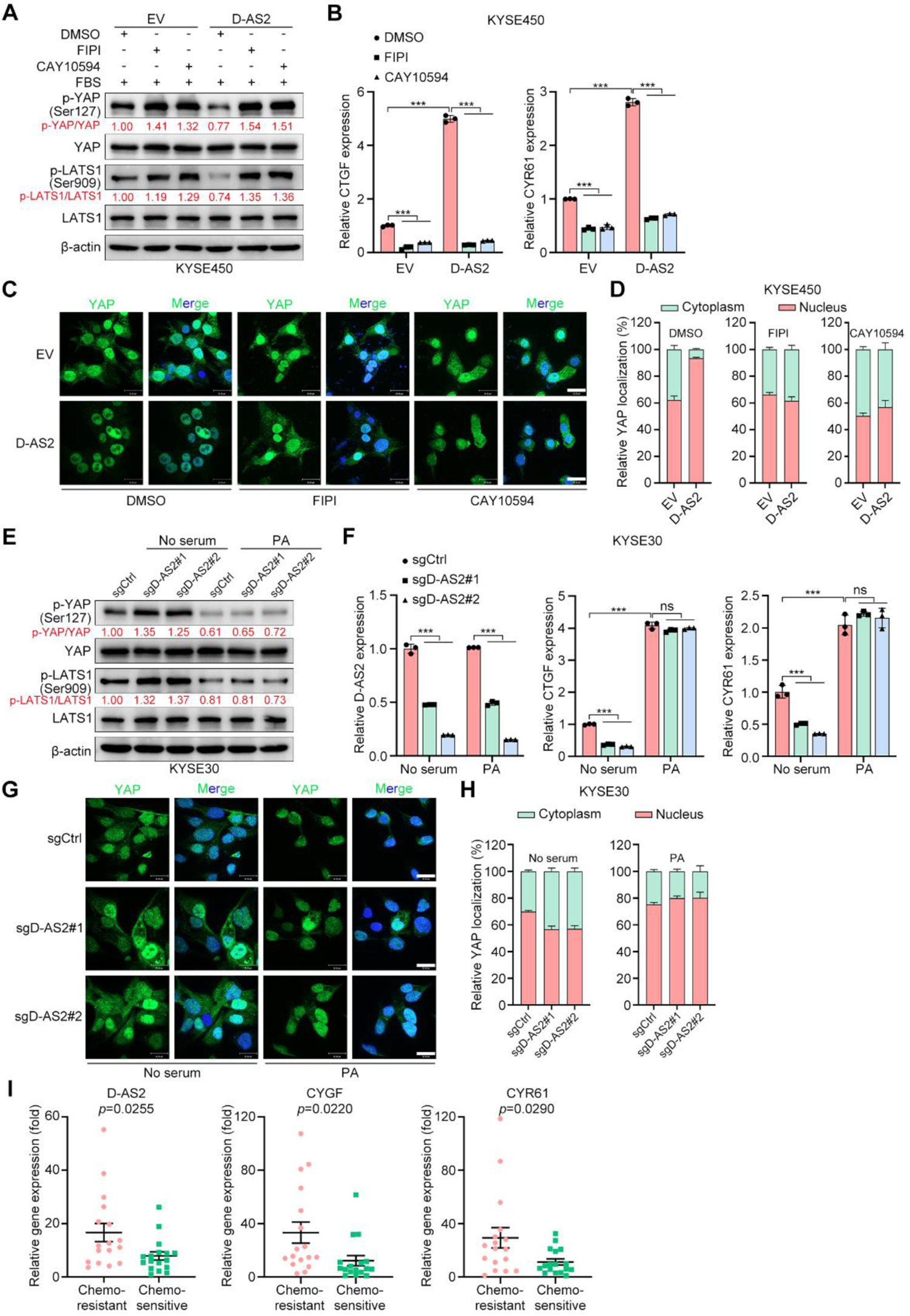
D-AS2 promotes PLD/PA-mediated YAP activation. (**A**) IB detection of YAP and LATS1 phosphorylation. KYSE450 cells expressing empty vector (EV) and D-AS2 were pretreated with FIPI (30 µM) or CAY10594 (20 µM) for 1 h. (**B**) RT-qPCR detection of the YAP target genes CTGF and CYR61. KYSE450 cells expressing EV and D-AS2 were pretreated with FIPI (30 µM) or CAY10594 (20 µM) for 1 h. The data are presented as the mean ± s.d. values; two-tailed *t* test, \*\*\**P* < 0.001; n = 3. (**C, D**) IF detection of YAP localization in KYSE450 cells expressing EV and D-AS2. Cells were pretreated with FIPI (30 µM) or CAY10594 (20 µM). YAP localization is shown in (C); scale bar, 30 µm. Cells from three different fields in three independent experiments were randomly selected for quantification of YAP localization (D). The data are expressed as the mean ± s.e.m. values. (**E**) IB detection of YAP and LATS1 phosphorylation. Serum-starved KYSE30 cells expressing control or D-AS2-targeting sgRNAs were pretreated with PA (100 µM) for 1 h. (**F**) RT-qPCR detection of the YAP target genes CTGF and CYR61. Serum-starved KYSE30 cells expressing control or D-AS2-targeting sgRNAs were pretreated with PA (100 µM) for 1 h. The data are presented as the mean ± s.d. values; two-tailed *t* test, \*\*\**P* < 0.001, ns: not significant; n = 3. (**G, H**) IF detection of YAP localization in KYSE30 cells expressing control and D-AS2-targeting sgRNAs and pretreated with PA (100 µM) for 1 h. YAP localization is shown in (G); scale bar, 30 µm. Cells from three different fields in three independent experiments were randomly selected for quantification of YAP localization (H). The data are expressed as the mean ± s.e.m. values. (**I**) RT-qPCR detection of the indicated genes expression in the samples with chemoresistance or chemosensitivity of esophageal SCC patients. The data are presented as the mean ± s.e.m. values; two-tailed *t* test; n = 17 for chemoresistant patients; n = 17 for chemosensitive patients.

**Figure 7.**
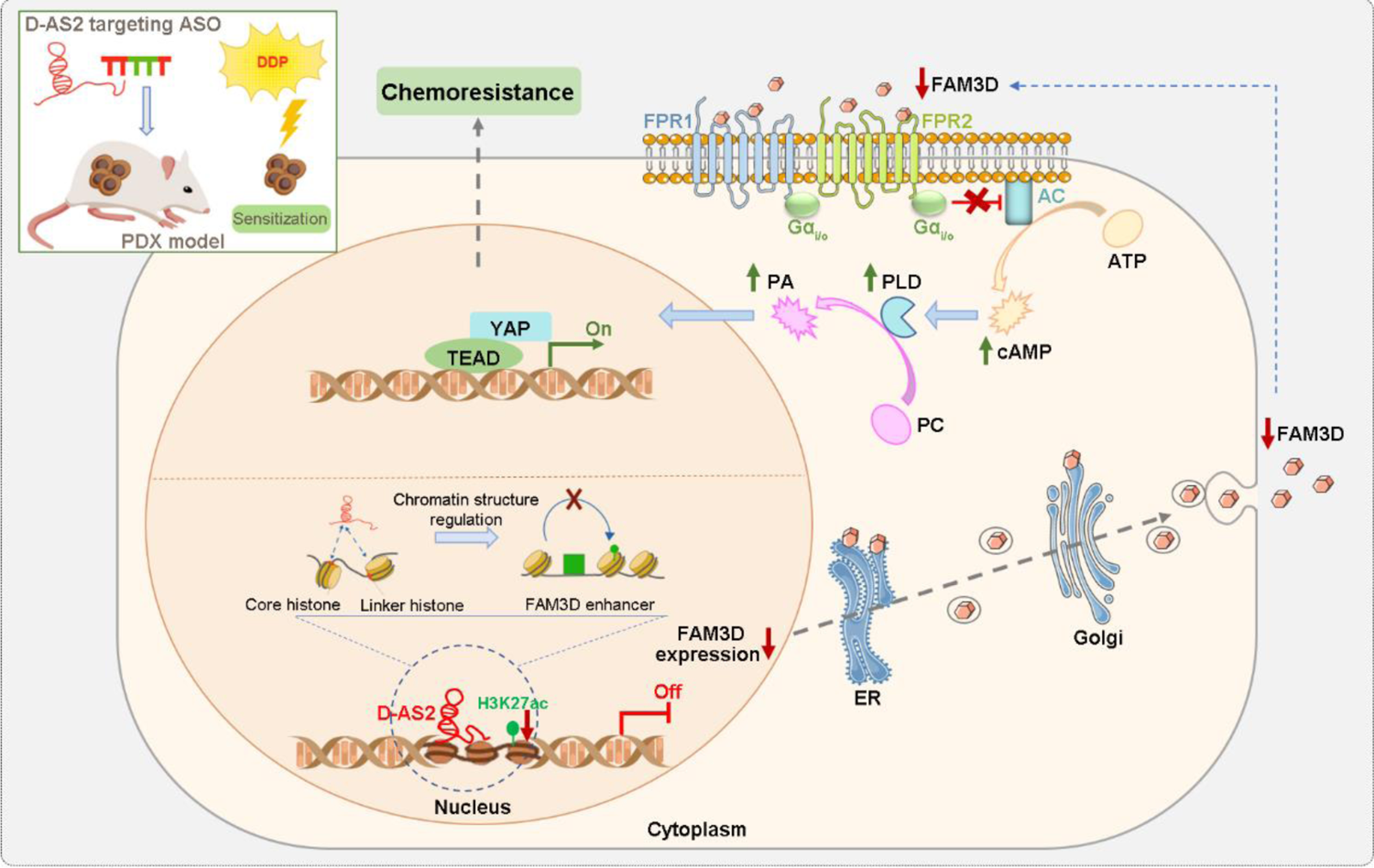
D-AS2 confers chemoresistance via regulating lipid metabolism and activating YAP signaling. D-AS2 associates with histones to regulate the distal elements of FAM3D and reduces extracellular FAM3D protein secretion. The reduction in extracellular FAM3D secretion activates PLD via FPR1 and FPR2 signaling and then increases PLD-dependent PA production. Furthermore, PA contributes to YAP nuclear translocation and activates nuclear YAP activity. ASO mediated D-AS2 targeting shows an encouraging effect on the sensitization of chemoresistant SCCs in *in vivo* studies.

The addition of PA was sufficient to reduce YAP and LATS1 phosphorylation and promote the transcription of the YAP downstream genes CTGF and CYR61 (Figs. 6E and 6F). In addition, depletion of D-AS2 led to sequestration of YAP in the cytoplasm, and PA treatment restored YAP nuclear translocation (Figs. 6G and 6H). As mentioned above, PA can be produced via three major metabolic pathways (Fig. S6H): PLD-catalyzed hydrolysis of PC to generate PA; LPAAT-catalyzed conversion of LPA to PA; and DGK-mediated phosphorylation of DAG to produce PA (22). To further determine whether D-AS2-mediated YAP activation is restricted to the PLD/PA pathway, we treated control and D-AS2-overexpressing KYSE450 and UM1 cells with CI 976 and R59-022, the specific inhibitors of LPAAT and DGK, respectively. Intriguingly, inhibition of either LPAAT by CI 976 or DGK by R59-022 failed to rescue YAP and LATS1 phosphorylation (Figs. S6I and S6J). At last, RT-qPCR was performed in SCC samples that exhibited either sensitivity or resistance during clinical treatment. The results indicated that D-AS2, CTGF and CYR61 were all significantly upregulated in the chemoresistant samples compared with in the chemosensitive samples (Fig. 6I), supporting that D-AS2 is a targetable regulator of YAP signaling and chemoresistance.

## Discussion

To overcome chemoresistance in SCC patients undergoing first-line treatment, we established two pairs of parental/chemoresistant cell models that mimic the clinical characteristics of chemoresistant cancer cells. We identified D-AS2, an antisense RNA of DLGAP1, as a critical lncRNA that contributes to the chemoresistance of SCC cells. A few very recent studies have shown that D-AS2 (47, 48) and another antisense RNA, DLGAP1-AS1 (49–51), play important roles in cancer. Despite the rapidly emerging studies, many aspects regarding the role of D-AS2 in cancer, including its role in chemoresistance, remain unclear. Our study provides strong evidence for the essential role of D-AS2 in SCCs and forms a basis for determining the other roles of D-AS2 or its function in other cancer types.

KEGG enrichment analysis of the differentially expressed mRNAs and the targets of the differentially expressed ncRNAs in the whole-transcriptome sequencing data set identified PLD signaling as the most significantly enriched specific metabolic pathway, and D-AS2 was identified as having the highest expression among the upregulated lncRNAs enriched in the PLD pathway. These findings suggest a potential relationship between D-AS2 and PLD. The PLD isoenzymes PLD1 and PLD2 are a major source of signaling-activated PA generation downstream of various cell surface receptors, including GPCRs, RTKs and integrins (52, 53). Numerous studies have reported PLD as a key modulator of cancer progression and chemoresistance (43, 54, 55). In particular, its catalytic product PA can activate YAP signaling (22, 23), and the activation of YAP signaling has been reported to play an oncogenic role in several contexts including chemoresistance (25, 26, 56, 57). Under Hippo pathway-activating conditions, mammalian sterile 20-like kinase 1/2 (MST1/2) are phosphorylated and activate LATS1/2 (58, 59). Phosphorylated LATS1 sequesters YAP in the cytoplasm by facilitating the phosphorylation of YAP at S127, leading to its degradation (60, 61). Herein, we propose a D-AS2/FAM3D-mediated PLD signaling pathway that increases PA production, promotes YAP activity, and contributes to chemoresistance in SCCs.

One surprising aspect of D-AS2 is that although its overexpression phenocopied the characteristics of the chemoresistant cell models, this overexpression exhibited a controversial oncogenic or tumor-suppressive effect. That is, D-AS2 promoted chemoresistance and suppressed cell proliferation. Cancer stem cells (CSCs) can undergo G0 arrest to avoid toxicity induced by chemotherapeutic agents and then induce relapse of cancer under appropriate conditions (62). Similarly, high expression of D-AS2 in SCC cells led to reduced proliferation caused by G0/G1 arrest. Furthermore, D-AS2 overexpression activated signaling through YAP, a factor recognized to mediate tumor stemness (26, 63). Therefore, one possible explanation for these dual effects is that D-AS2 promotes CSC-like transformation. These results suggest that targeting cancer cells with high D-AS2 expression may be an effective strategy to eliminate the CSC pool and prevent cancer relapse.

To further identify the critical downstream targets of D-AS2 that promote chemoresistance and regulate lipid metabolism, we analyzed the differentially expressed genes of D-AS2-overexpressing cells and two chemoresistant cell models and found that FAM3D, an extracellular secreted protein, mediates the function of D-AS2 in chemoresistance. The FAM3 gene family contains 4 members: FAM3A, FAM3B, FAM3C, and FAM3D. The first three members have been suggested to be involved in regulating glucose and lipid metabolism (64), while the role of FAM3D in metabolism remains largely unknown. Our ATAC-seq results indicated that D-AS2 reduced the enrichment of the transcriptional activation marker H3K27ac in the FAM3D enhancer region, in turn reducing the extracellular secretion of FAM3D. FAM3D has been reported to act as a dual agonist of FPR1 and FPR2 to induce Mac-1-mediated neutrophil recruitment and to aggravate abdominal aortic aneurysm (AAA) development through FPR-related Gi protein and β-arrestin signaling (39, 65). Herein, we found that FAM3D, as a specific ligand of FPR1 and FPR2, activated FPR Gα_i_ signaling in SCCs, resulting in decreased PLD activity. On the other hand, the D-AS2-mediated decrease in extracellular FAM3D secretion activated PLD and increased PA production through FPR1 and FPR2 signaling, resulting in activation of YAP signaling and chemoresistance.

## Author contributions

Y.N. and Q.L. designed the experiments. Y.N., Q.L., and X.W. performed most of the experiments and data analysis. S.L., P.Z., and W.C. assisted with animal studies. A.Z. helped collect clinical data. Y.N. drafted the manuscript with assistance from Q.L., X.W., and Z.L. All authors reviewed the manuscript. Z.L. organized and supervised this study.

## Acknowledgments

We are grateful to Dr. Yutaka Shimada (Kyoto University, Japan) for the esophageal SCC cell lines. We thank Dr. Yin Li for the SCC samples. We appreciate Beijing Qinglian Biotech Co., Ltd., for performing the MS analysis. We thank Yong Peng and Lingsheng Chen (Shanghai Applied Protein Technology Co., Ltd., China) for performing the lipidomic analyses.

