## Supplemental figures and tables for "DLGAP1-AS2-Mediated Phosphatidic Acid Synthesis Confers Chemoresistance via Activation of YAP Signaling"

### Supplementary figures

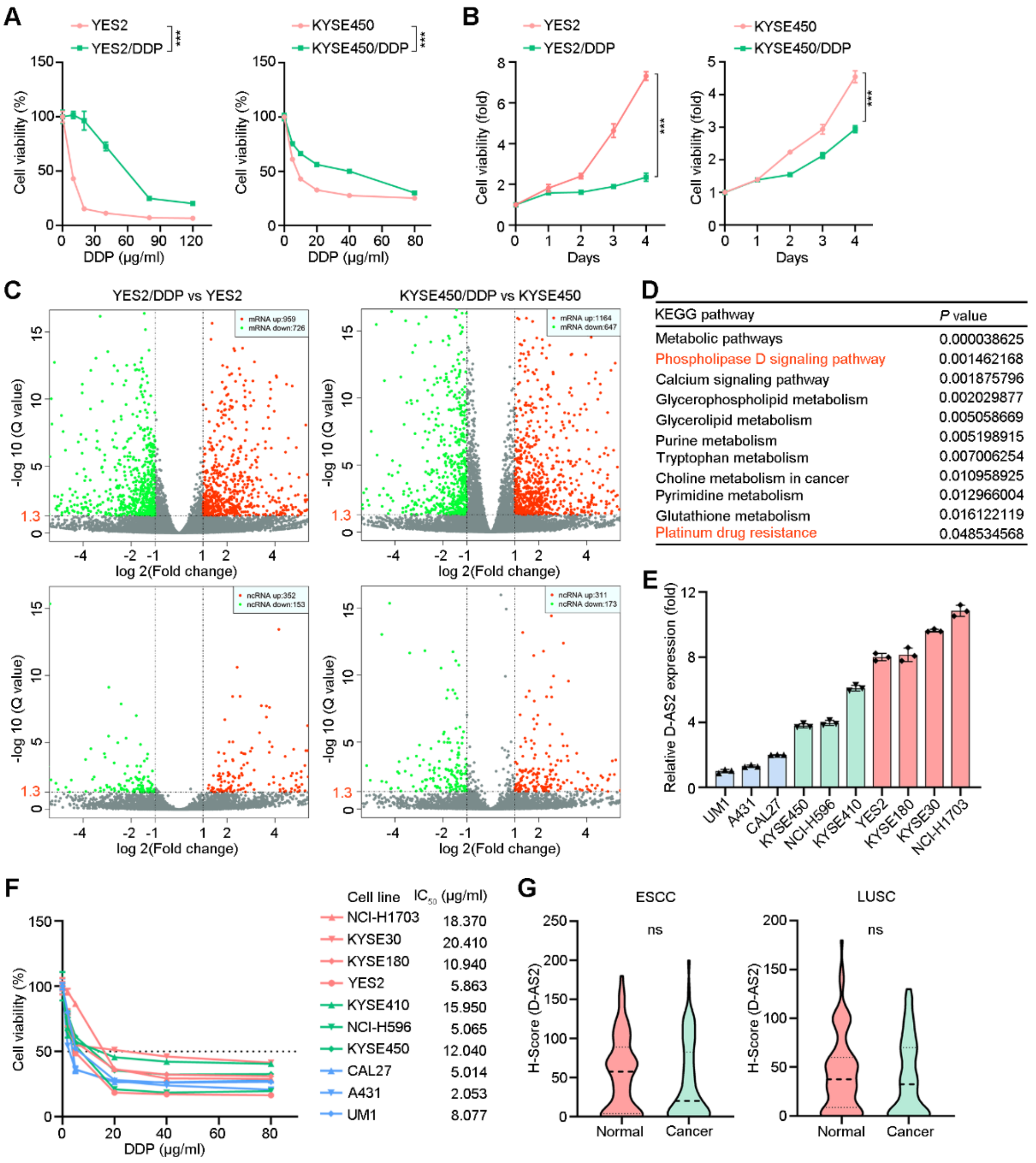


**Figure S1. D-AS2 shows increased expression in chemoresistant SCC cells.**

(**A**) Relative viability of parental and chemoresistant SCC cells after DDP treatment for 24 h. The data are presented as the mean ± s.d. values; two-way ANOVA, ***$P < 0.001$; n = 5. (**B**) In vitro growth curves of parental and chemoresistant SCC cells. The data are presented as the mean ± s.d. values; two-tailed $t$ test, ***$P < 0.001$; n = 5. (**C**) Volcano plots showing upregulated and downregulated mRNAs and ncRNAs in chemoresistant cells compared to the corresponding parental cells. (**D**) KEGG enrichment analysis of the predicted target genes downstream of the 38 ncRNAs upregulated and downregulated in both chemoresistant cell lines. (**E**) RT–qPCR detection of D-AS2 expression in SCC cell lines. The data are presented as the mean ± s.d. values; n = 3. (**F**) Relative cell viability and DDP $IC_{50}$ values in SCC cell lines after DDP treatment for 24 h. The data are presented as the mean ± s.d. values; n = 5. (**G**) Statistical analysis of D-AS2 expression in the two cohorts of esophageal and lung SCC tissues and adjacent normal tissues. The data are presented as the mean ± s.e.m. values; two-tailed *t* test, ns: not significant; n = 70 for esophageal SCC; n = 90 for lung SCC.

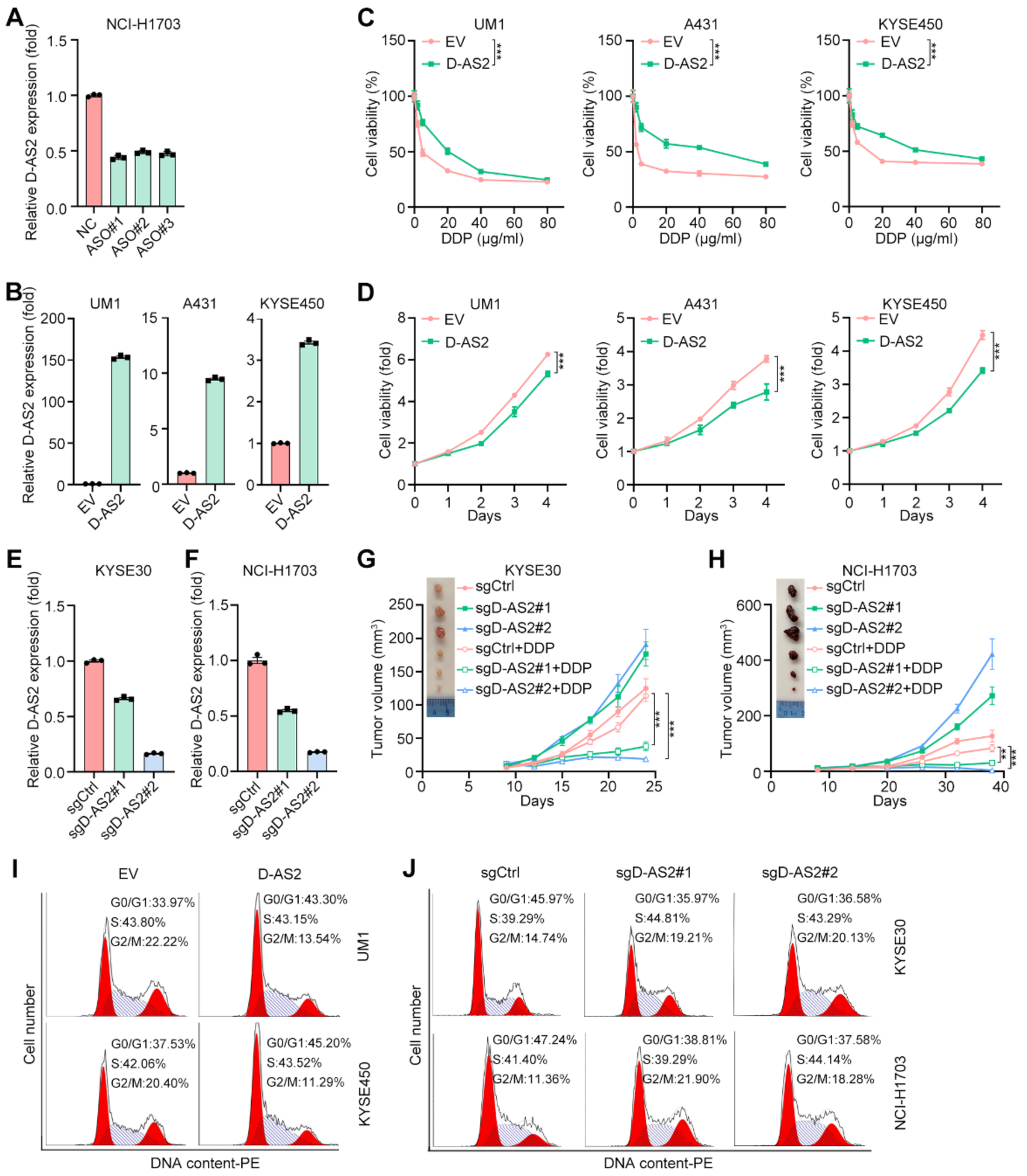

**Figure S2. D-AS2 suppresses SCC cell proliferation but leads to chemoresistance.** (**A**) RT–qPCR detection of D-AS2 expression in NCI-H1703 cells transfected with negative control (NC) or D-AS2 (ASO#1, ASO#2, and ASO#3) ASOs. The data are presented as the mean ± s.d. values; n = 3. (**B**) RT–qPCR detection of D-AS2 expression in SCC cells expressing empty vector (EV) or D-AS2. The data are presented as the mean ± s.d. values; n = 3. (**C**) Relative viability of SCC cells expressing EV or D-AS2 after DDP treatment for 24 h. The data are presented as the mean ± s.d. values; two-way ANOVA, ****P* < 0.001; n = 5. (**D**) In vitro growth curve of SCC cells expressing EV or D-AS2. The data are presented as the mean ± s.d. values; two-tailed *t* test, ***$P$ < 0.001; n = 5. (**E, F**) RT–qPCR detection of D-AS2 expression in KYSE30 and NCI-H1703 cells expressing control or D-AS2-targeting sgRNAs. The data are presented as the mean ± s.d. values; n = 3. (**G, H**) Representative images and volumes of xenografts derived from KYSE30 and NCI-H1703 cells expressing control or D-AS2-targeting sgRNAs. The data are presented as the mean ± s.e.m. values; two-tailed *t* test, ***$P$ < 0.001, **$P$ < 0.01; n = 8. (**I, J**) Representative images from the cell cycle analysis of the indicated cells.

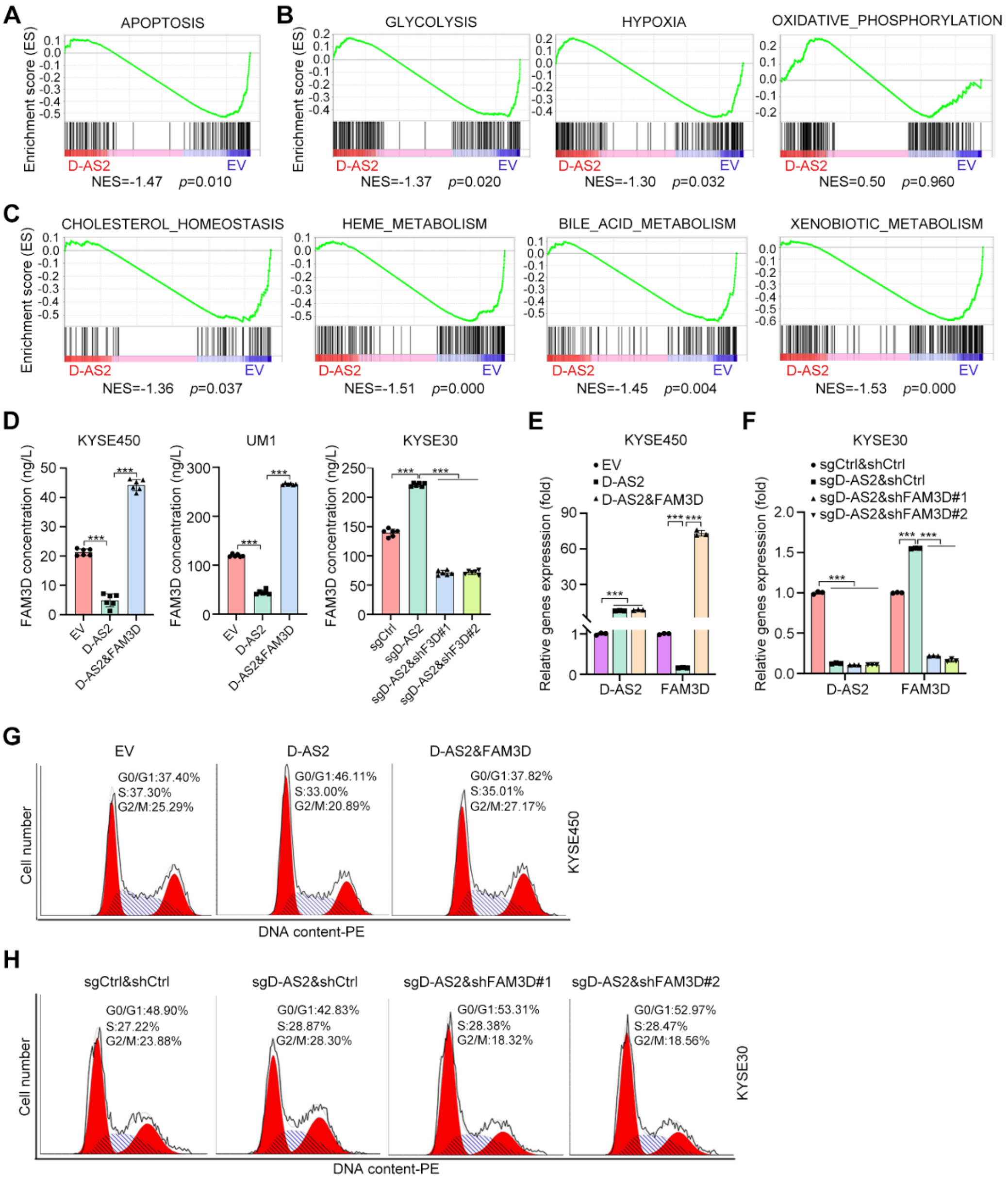


**Figure S3. FAM3D mediates the function of D-AS2.**

(**A-C**) GSEA of differentially expressed mRNAs in D-AS2-overexpressing KYSE450 cells compared to control cells. (**D**) ELISA of the FAM3D concentration in the indicated cells. The data are presented as the mean ± s.d. values; two-tailed *t* test, $^{***}P < 0.001$; n = 6. (**E, F**) RT–qPCR detection of D-AS2 and FAM3D expression in the indicated cells. The data are presented as the mean ± s.d. values; two-tailed *t* test, $^{***}P < 0.001$; n = 3. (**G, H**) Representative images from the cell cycle analysis of the indicated cells.

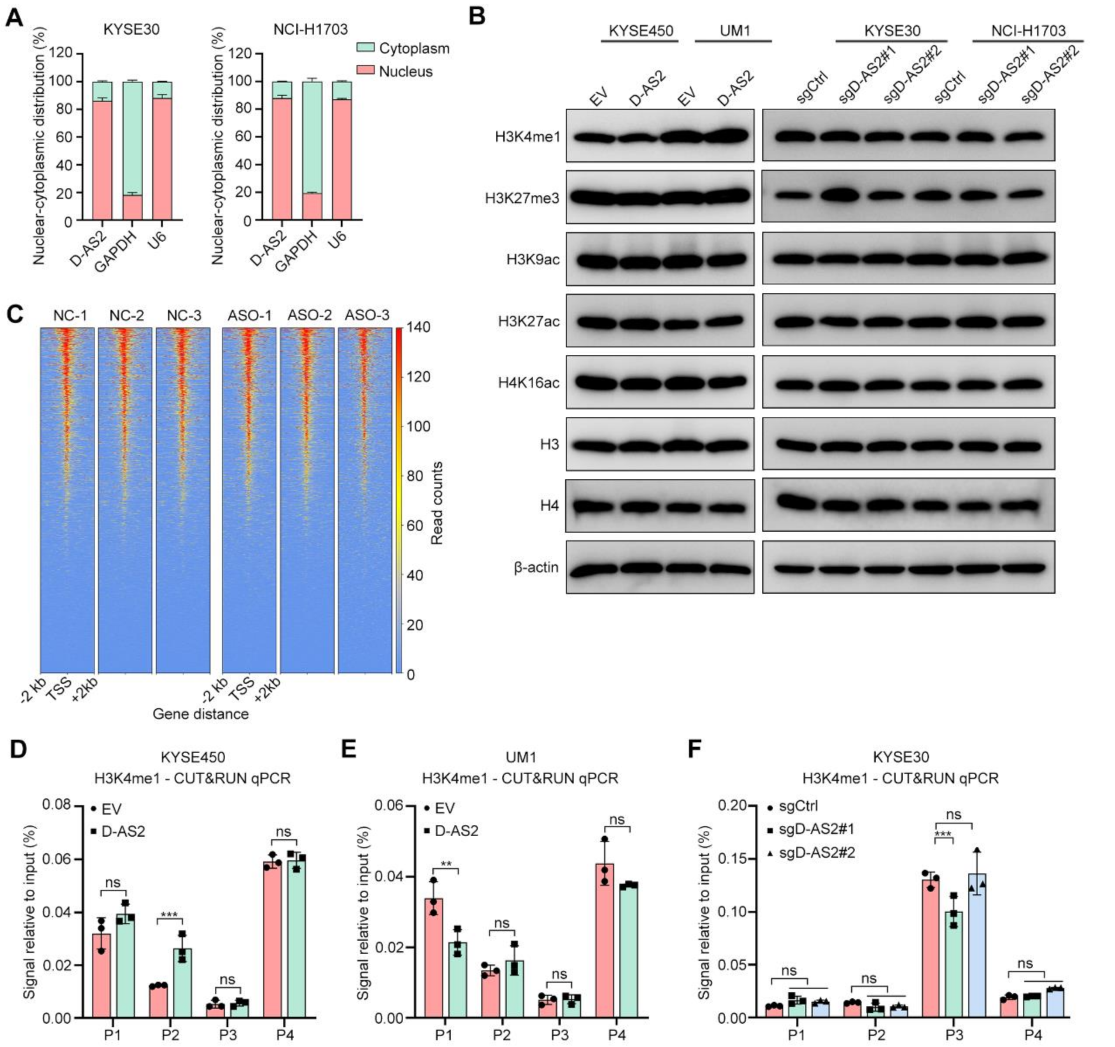


**Figure S4. D-AS2 primarily regulates gene expression via distal elements.**

(**A**) RT–qPCR detection of D-AS2 expression in the cytoplasmic and nuclear fractions. (**B**) IB detection of histone marks in D-AS2-depleted and D-AS2-overexpressing SCC cells. (**C**) Heatmaps showing ATAC signals within the region ±2 kb around the TSS. (**D-F**) CUT&RUN assay of the FAM3D enhancer region in D-AS2-depleted and D-AS2-overexpressing SCC cells with antibodies against H3K4me1. The data are presented as the mean ± s.d. values; two-tailed *t* test, ***$P < 0.001$, **$P < 0.01$, ns: not significant; n = 3.

**Figure S5. D-AS2 activates PLD through FAM3D-mediated FPR1 and FPR2 signaling.**

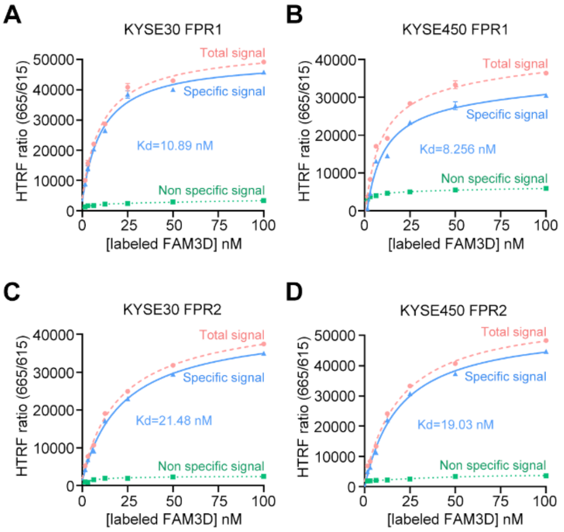


(**A-D**) Saturation curve of FAM3D binding to FPR1 and FPR2. KYSE30 and KYSE450 cells transiently expressing FPR1- and FPR2-tagged SNAP were incubated with increasing concentrations of labeled FAM3D. A prominent homogeneous time-resolved fluorescence (HTRF) signal is visible. Nonspecific binding was measured by adding 10 μM FPR Agonist 43 to the wells. The dissociation constants ($K_d$ values) are shown. The data are presented as the mean ± s.d. values; n = 3.

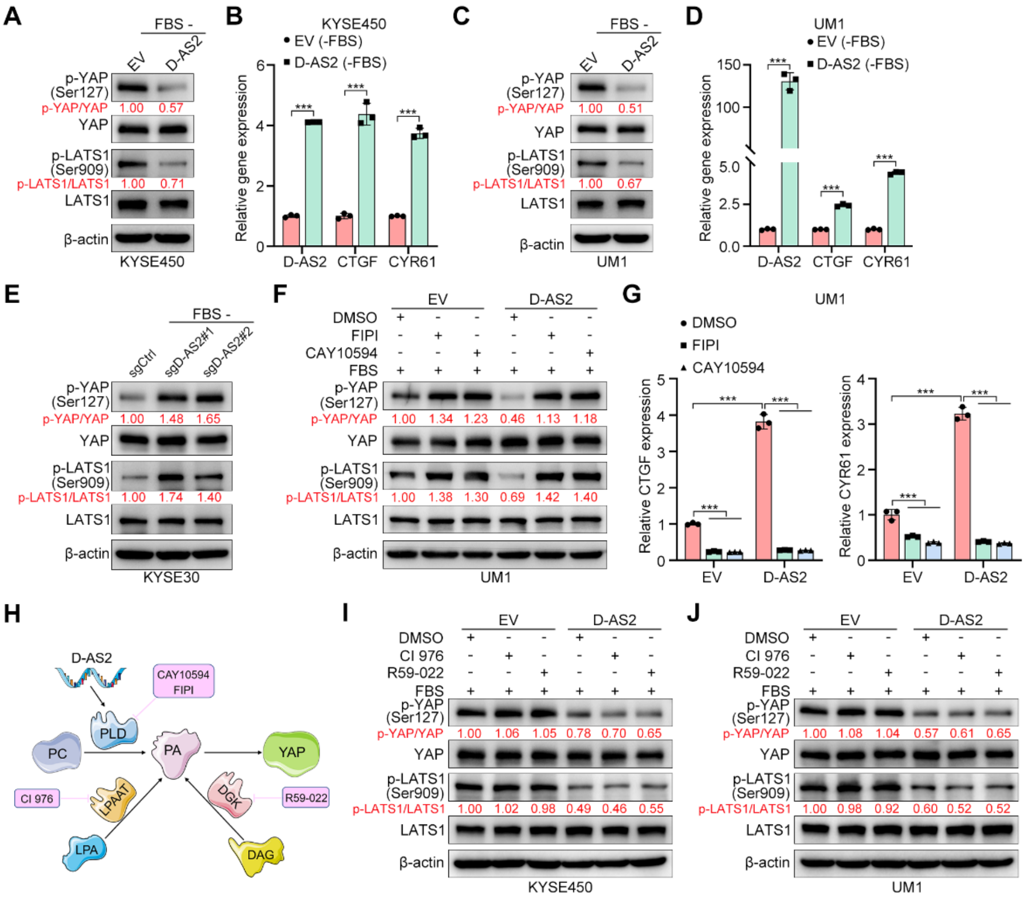


**Figure S6. D-AS2 activates YAP signaling through PLD/PA.**

(**A**) IB detection of phosphorylated YAP and LATS1 in serum-starved control and D-AS2-overexpressing KYSE450 cells (**B**) RT–qPCR detection of the YAP target genes CTGF and CYR61 in serum-starved control and D-AS2-overexpressing KYSE450 cells. The data are presented as the mean ± s.d. values; two-tailed *t* test, ****P* < 0.001; n = 3. (**C**) IB detection of phosphorylated YAP and LATS1 in serum-starved control and D-AS2-overexpressing UM1 cells (**D**) RT–qPCR detection of the YAP target genes CTGF and CYR61 in serum-starved control and D-AS2-overexpressing UM1 cells. The data are presented as the mean ± s.d. values; two-tailed *t* test, ****P* < 0.001; n = 3. (**E**) IB detection of phosphorylated YAP and LATS1 in serum-starved control and D-AS2-depleted KYSE30 cells. (**F**) IB detection of YAP and LATS1 phosphorylation. UM1 cells expressing empty vector (EV) and D-AS2 were treated with FIPI (30 μM) or CAY10594 (20 μM) for 1 h. (**G**) RT–qPCR detection of the YAP target genes CTGF and CYR61. UM1 cells expressing EV and D-AS2 were treated with FIPI (30 μM) or CAY10594 (20 μM) for 1 h. The data are presented as the mean ± s.d. values; two-tailed *t* test, ***$P < 0.001$; n = 3. (**H**) Schematic illustration of the three metabolic pathways for PA production. The inhibitors targeting each pathway or enzyme are indicated. (**I, J**) IB detection of phosphorylated YAP and LATS1 in control and D-AS2-overexpressing KYSE450 and UM1 cells. Cells were pretreated with CI 976 (20 μM) or R59-022 (20 μM) for 30 min.

**Supplementary tables**

**Table S1. D-AS2 probe sequences used for ISH**

| Sequence (5’ to 3’) | Name |
| --- | --- |
| AATGGTATTTCCATTTATATAAGAAGCGCCTAAGAAATGC | Probe#1 |
| AATTTGATGCCAACTTTATGTGTAAAGAAGCTAACTCCTG | Probe#2 |
| AAGAAGAAACTGAATTTGAAGTGGATTCTTACAAAGGAAA | Probe#3 |

**Table S2. RT–qPCR primers**

| Sequence (5' to 3') | Name |
|---|---|
| CTCGCTTCGGCAGCACA | U6-F |
| AACGCTTCACGAATTTGCGT | U6-R |
| CCGGGAAACTGTGGCGTGATGG | GAPDH-F |
| AGGTGGAGGAGTGGGTGTCGCTGTT | GAPDH-R |
| GCGCCTAAGAAATGCCTGT | DLGAP1-AS2-F |
| AGCTGTTCATTCAGCCACGA | DLGAP1-AS2-R |
| CTGCCCAGCCAACTACTTTG | FAM3D-F |
| CTCCCGTGGTTCCATTCAC | FAM3D-R |
| CCAATGACAACGCCTCCTG | CTGF-F |
| TGGTGCAGCCAGAAAGCTC | CTGF-R |
| AGCCTCGCATCCTATACAACC | CYR61-F |
| TTCTTTCACAAGGCGGCACTC | CYR61-R |

**Table S3. shRNA and sgRNA sequences**

| Sequence (5’ to 3’) | Name |
|---|---|
| GCACCTAGTGAAATTCCTTAA | shFAM3D#1 |
| GCCCAGACACAAACAAATACG | shFAM3D#2 |
| CACCGGCACGCTCTCTGACAGCATC | sgDLGAP1-AS2#1 |
| CACCGGAACGTCACAGGCATTTCTT | sgDLGAP1-AS2#2 |

**Table S4. CUT&RUN qPCR primers targeting the FAM3D enhancer region**

| Sequence (5' to 3') | Name |
|---|---|
| TGGAAGGGATGTTGGGCAGTGA | Primer1-F |
| TAGCGAGAGGCAGCAGAGTGAA | Primer1-R |
| ACCTCCAGGGCACGATTCCTATAC | Primer2-F |
| CAGGCAACAGGAAGTGCTGACAT | Primer2-R |
| ACCCTGGTGGAGAAAGAAGTGACTAT | Primer3-F |
| CTGACTGTGAAGGCTGACAAGGTG | Primer3-R |
| ACCTTGTCAGCCTTCACAGTCAG | Primer4-F |
| GAAACCCACTTCCTTCCTTCCAACA | Primer4-R |
